# Non-invasive temporal interference electrical stimulation of the human hippocampus

**DOI:** 10.1101/2022.09.14.507625

**Authors:** Ines R. Violante, Ketevan Alania, Antonino M. Cassarà, Esra Neufeld, Emma Acerbo, Romain Carron, Adam Williamson, Danielle L. Kurtin, Edward Rhodes, Adam Hampshire, Niels Kuster, Edward S. Boyden, Alvaro Pascual-Leone, Nir Grossman

## Abstract

Deep brain stimulation (DBS) via implanted electrodes is used worldwide to treat patients with severe neurological and psychiatric disorders however its invasiveness precludes widespread clinical use and deployment in research. Temporal interference (TI) is a strategy for non-invasive steerable DBS using multiple kHz-range electric fields with a difference frequency within the range of neural activity. Here we report the validation of the non-invasive DBS concept in humans. We used electric field modelling and measurements in a human cadaver to verify that the locus of the transcranial TI stimulation can be steerably focused in the hippocampus with minimal exposure to the overlying cortex. We then used functional magnetic resonance imaging (fMRI) and behaviour experiments to show that TI stimulation can focally modulate hippocampal activity and enhance the accuracy of episodic memories in healthy humans. Our results demonstrate targeted, non-invasive electrical stimulation of deep structures in the human brain.

## INTRODUCTION

A multitude of brain disorders have debilitating impacts on quality of life, with neurological conditions increasingly recognised as major causes of death and disability, accounting for approximately 30% of the global burden of disease (GBD)^1^. Most patients with brain disorders are unamenable to any form of pharmacological treatment^2,3^. Physical means of brain stimulation, known as ‘neuromodulation’, represent a tenable, non-pharmacological treatment strategy that acts through direct control of the aberrant neural activity underpinning the diseases or their symptomatic manifestation. Invasive electrical deep brain stimulation (DBS) has been used worldwide to treat patients with severe movement disorders, such as Parkinson’s disease^4^, and affective disorders, such as obsessive-compulsive disorder^5^. In addition, DBS is being investigated as a treatment for other conditions, such as depression^6,7^ and Alzheimer’s disease^8,9^. However, the risks associated with brain surgery make exploration of different brain targets difficult and limits potential therapeutic impact^4,10^.

Non-invasive stimulation methods, such as transcranial magnetic stimulation (TMS) and transcranial electrical stimulation (TES), have been used in many human clinical investigations^11,12^. However, their ability to directly stimulate deeper brain structures is achieved at the expense of inducing stronger stimulation of the overlying cortical areas, resulting in unanticipated side effects that can approach the limits of safety guidelines^13^.

We recently reported a strategy for sculpting electrical fields to enable focused yet non-invasive neural stimulation at depth^14^. By delivering multiple electric fields to the brain at slightly different frequencies within the kHz range—which are themselves higher than the frequency range of endogenous neural activity but for which the difference frequency is sufficiently low to drive neural activity—neurons can be electrically activated at a selected focus without driving neighbouring or overlying regions. We call this strategy temporal interference (TI) stimulation, since the interference of multiple electric fields is what enables its focality: neural stimulation will occur at the targeted region, for which the amplitude of the electric field envelope modulation, at the difference frequency, is of greater magnitude (**Fig. 1a)**. We validated the TI stimulation concept in the mouse brain both electrophysiologically and by imaging *c-fos* as a marker of neural activation. We demonstrated that TI stimulation can selectively mediate activation in a deep neural structure, i.e., the hippocampus, without activating the overlying cortical neurons, and steerably target brain regions without physically moving the electrodes^14^.

**Fig. 1:**
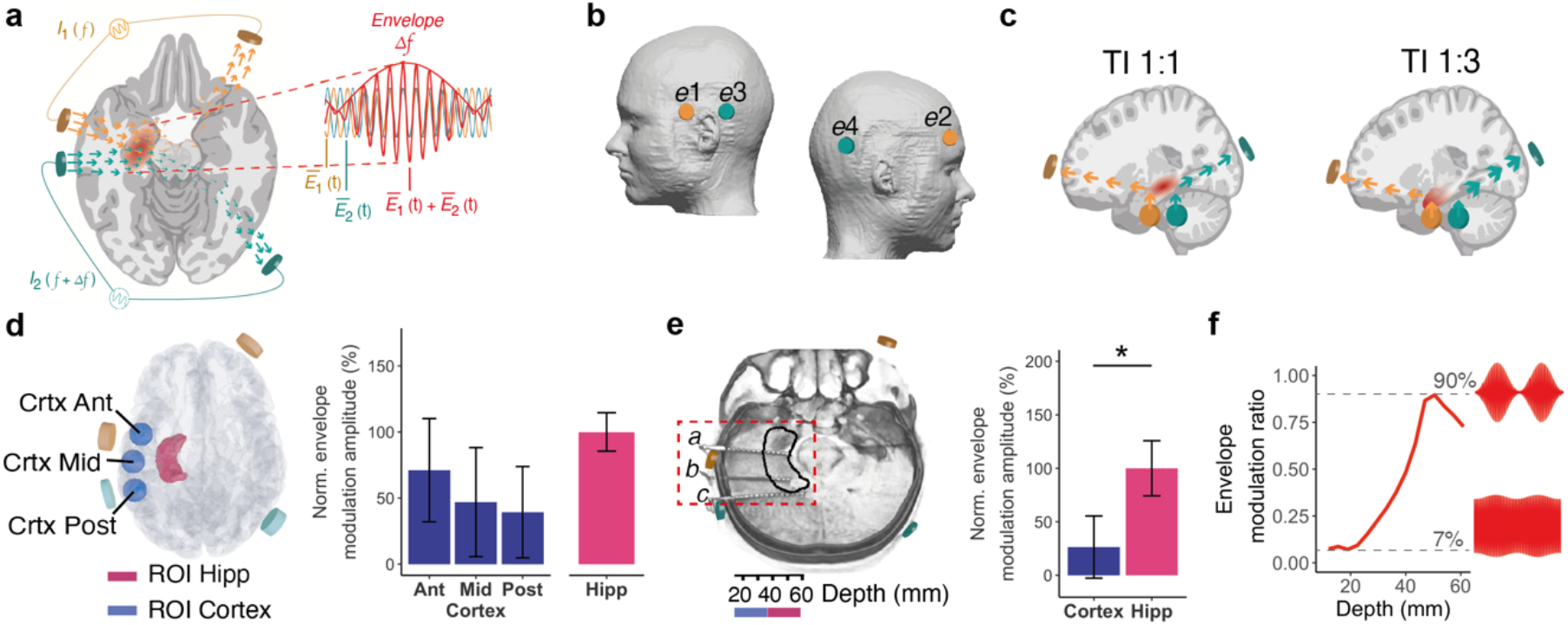
Fundamentals of Temporal Interference (TI) hippocampal stimulation and validation using computational modelling and cadaver measurements. **a-c**, Concept of transcranial TI hippocampal stimulation. **a**, Two current sources *I*_*1*_ and *I*_*2*_ are applied simultaneously via electrically isolated pairs of scalp electrodes (orange and green) at kHz frequencies *f*1 and *f*2, with a small frequency difference f = *f*1-*f*2 within the range of neural activity. The applied currents generate oscillating electric fields Ē_1_(t) and Ē_2_(t) inside the brain (orange and green arrows, respectively). Superposition of these fields, Ē_1_(t) + Ē_2_(t), results in an envelope amplitude that is modulated periodically at Δf. The peak amplitude of the envelope modulation can be localised in deep brain structures such as the hippocampus (highlighted in red). **b**, Schematic of electrode configuration targeting the left hippocampus displayed in the standard head model. Electrodes *e*1 and *e*2 formed one electrode pair (orange) and electrodes *e*2 and *e*4 a second electrode pair (green), corresponding to current sources *I*_*1*_ and *I*_*2*_ in A. Electrodes *e*1 and *e*3 at the narrow base of the trapezoid were located at nasion plane of the left hemisphere, symmetrically above the anterior-posterior midline of the hippocampus, with a 5 cm distance between the electrode centres. Electrodes *e*2 and *e*4 at the wider base of the trapezoid were located at a plane above the eyebrow on the right hemisphere with approximately 16 cm distance between the electrode centres. All electrodes were 1.5 cm x 1.5 cm square with rounded corners. **c**, Illustration of how tuning the current ratios steers the TI stimulation locus along the hippocampal longitudinal axis. TI stimulation with 1:1 current ratio (‘TI 1:1’) and stimulation locus in the middle region, left panel; TI stimulation with 1:3 current ratio (‘TI 1:3’) and locus in the anterior region, right panel. By reducing the current amplitude in one electrode pair and increasing the current amplitude in the second electrode pair by the same amount (i.e., keeping the current sum fixed), the stimulation locus can be steered towards the electrode pair with the smaller current amplitude^15^. **d**, Computation of TI stimulation locus in a human anatomical model. Fields in regions of interest (ROIs) in the left (stimulated) hippocampus and its overlying cortex, showing schematic of the regions of interest (ROIs); Ant – anterior, Mid – middle, Post – posterior, left panel; fields’ envelope modulation amplitude in the ROIs, right panel; values are median∓standard deviation (SD) normalised to the hippocampal value. For the fields’ absolute amplitude see Fig. S1. **e-f**. Measurement of TI stimulation locus in a human cadaver with *I*_*1*_ (2 kHz, 1mA) and *I*_*2*_ (2.005 kHz, 1mA). **e**, CT image of the human cadaver with intracranial electrode leads *a, b* and *c* implanted in the left mesial temporal lobe. Each electrode consisted of 15 electrode contacts; Black contour, approximate location of the left hippocampus; orange and green stimulation electrodes, left panel. Right panel: Amplitudes of the envelope modulation in the left (stimulated) hippocampus and its overlying cortex showing higher envelope amplitude at the hippocampus; shown values are median±SD averaged across electrodes *a*-*c* and then normalised to hippocampal value. *p<0.05, See **Table S1** for full statistics and **Fig. S2** for additional amplitude maps. **f**, Envelope modulation ratio vs depth for electrode *b*, showing increasing envelope modulation with depth.

Here, we aim to test the translation of these results by investigating the application of TI to the human hippocampus. Earlier human studies tested TI stimulation of cortical structures^15-17^, but the crucial non-invasive DBS capability has not been validated to date. We first focused on validating the locus of TI stimulation using computational modelling and cadaver measurements. We followed this by performing simultaneous TI and functional magnetic resonance imaging (fMRI) experiments designed to explore physiological changes in brain activity in response to stimulation and provide evidence for target engagement. Finally, we tested the behavioural impact of delivering TI stimulation to the hippocampus in healthy participants. We demonstrate the safety and tolerability of TI in humans, the ability to focally target the stimulation locus to the hippocampus, and the capacity to modulate hippocampal activity and behavioural performance.

## RESULTS

### Validation of hippocampal targeting

We first examined whether the locus of TI stimulation can be localised to the hippocampus with minimal exposure of the overlying cortex. We positioned two pairs of electrodes on the scalp in a configuration that targets the left hippocampus (**Fig. 1b**) and computed the field distribution in the established anatomical MIDA model^18^, that distinguishes a large number of tissue classes derived from high-resolution multi-modal MRI data, and accounts for electrical conductivity anisotropy and neural orientation based on diffusion tensor imaging (DTI). We applied two sinusoidal currents at 2.005 kHz and 2 kHz (resulting in a Δf envelope frequency of 5 Hz) and an equal amplitude of 1 mA (current density ∼0.45 mA/cm^2^), i.e., TI with 1:1 current ratio (‘TI 1:1’, **Fig. 1c** left), and computed the fields’ envelope modulation amplitude and absolute amplitude along the (DTI-derived) principal fibres axis in regions-of-interest (ROIs) at the left, i.e. stimulated, hippocampus (‘Hipp’) and the overlying cortical regions, both underneath the stimulation electrodes (anterior, ‘Crtx_Ant’; posterior, ‘Crtx_Post’) and between the stimulation electrodes (middle, ‘Crtx_Mid’), (**Fig. 1d**). The fields’ envelope modulation amplitude in the hippocampus was 30-60% larger than in the overlying cortical regions (Hipp, 0.26±0.04 V/m median∓SD; Crtx_Ant, 0.18∓0.10; Crtx_Mid, 0.12∓0.11; Crtx_Post, 0.10∓0.09; **Fig. 1d** and **Fig. S1**). In contrast, the fields’ absolute amplitude in the hippocampus was smaller than in the overlying cortical region underneath the Ant electrode (Hipp, 0.29∓0.04 V/m; Crtx_Ant, 0.30∓0.17; Crtx_Mid, 0.23∓0.11; Crtx_Post, 0.21∓0.13, **Fig. S1**).

Given the distinctive functional organization along the hippocampal longitudinal axis^19^, we next explored the relative distribution of the TI electric fields between the ‘Ant’, ‘Mid’ and ‘Post’ regions of the hippocampus. The model showed similar envelope modulation amplitudes across hippocampal regions relative to total hippocampal exposure (**Fig. S1**). Since the anterior hippocampus has been explicitly implicated in successful associative encoding^20^, we explored whether the locus of the TI electric fields can be steered anteriorly. By reducing the current in the anterior electrode pair *e1*-*e2* to 0.5 mA (∼0.225 mA/cm^2^) and increasing the amplitude in the posterior electrode pair *e3*-*e4* to 1.5 mA (∼0.675 mA/cm^2^), i.e., TI with 1:3 current ratio (‘TI 1:3’, **Fig. 1c** right), we found that TI 1:3 stimulation could increase the relative envelope modulation amplitude in the Ant hippocampal region (**Fig. S1**).

To validate that the locus of TI stimulation could indeed be targeted to the hippocampus, we applied the same electrode configuration and sinusoidal currents (**Fig. 1c**, TI 1:1) to a human cadaver and measured the electrical potential using intracranial electrodes implanted in the left mesial temporal lobe, perpendicular to the hippocampus long axis (**Fig. 1e**). Consistent with our modelling, we found that the normalised envelope modulation amplitude was ∼75% larger in the hippocampus compared to the overlying cortex (**Fig. 1e** and **Table S1**). The largest electric field’s envelope modulation amplitude along the recording electrode *b*, between the two stimulation electrodes *e1* and *e2* (i.e., a field direction perpendicular to the hippocampal longitudinal axis), was ∼0.1 V/m at a depth of ∼44 mm (consistent with the location of the hippocampus^21^) per 1 mA applied current (current density ∼0.45 mA/cm^2^). The envelope modulation ratio along electrode *b* was low at the cortex (∼7% at 12 mm depth) and high near the hippocampus (∼90% at 50 mm depth, **Fig. 1f**). In contrast, the absolute amplitude was largest in the overlying cortical region, the normalised amplitude was ∼50% larger in the cortex compared to the underlying hippocampus (**Table S1**). The distribution of the absolute amplitude was similar when we applied two sinusoids at the Δf frequency of 5 Hz (**Fig. S2**).

### Probing the physiological effect of TI stimulation

After establishing that the TI stimulation locus could be focally and steerably targeted to the hippocampus, we aimed to test whether the stimulating fields could modulate hippocampal neural activity. We applied TI stimulation to the hippocampus of twenty healthy participants (mean age 27.1 ± 7.6 SD, 11 females) while measuring brain activity using blood-oxygenation-level-dependent (BOLD) fMRI. To evoke hippocampal activity, the stimulation was delivered while the participants performed a hippocampal-dependent face-name paired associative task (**Fig. 2a**), known to robustly evoke BOLD signal in the hippocampus^22,23^. Stimulation was applied using the same electrode configuration as before (**Fig. 1b)** across three conditions: 1) TI 1:1 (2.005 kHz, 2 mA, 0.9 mA/cm^2^; and 2 kHz, 2 mA, 0.9 mA/cm^2^), and 2) TI 1:3 (2.005 kHz, 1 mA, 0.45 mA/cm^2^; and 2 kHz, 3 mA, 1.35 mA/cm^2^), but with two-fold larger amplitudes, and 3) a sham condition (2.005 kHz, 0 mA, 0 mA/cm^2^; and 2 kHz, 0 mA, 0 mA/cm^2^). We chose a Δf difference frequency of 5 Hz within the theta-band due to the evidential bases for the role of hippocampal theta-band oscillation in episodic memory^24-26^. Stimulation was only applied for a short period during the encoding period of each block (i.e., 32 s). Each participant received three blocks of each stimulation condition (i.e., a total of 96 s per stimulation condition) in a counterbalanced order between participants (**Fig. 2a**). This ON/OFF design to minimise build-up while maximising signal-to-noise-ratio to assess physiological responses.

**Fig. 2:**
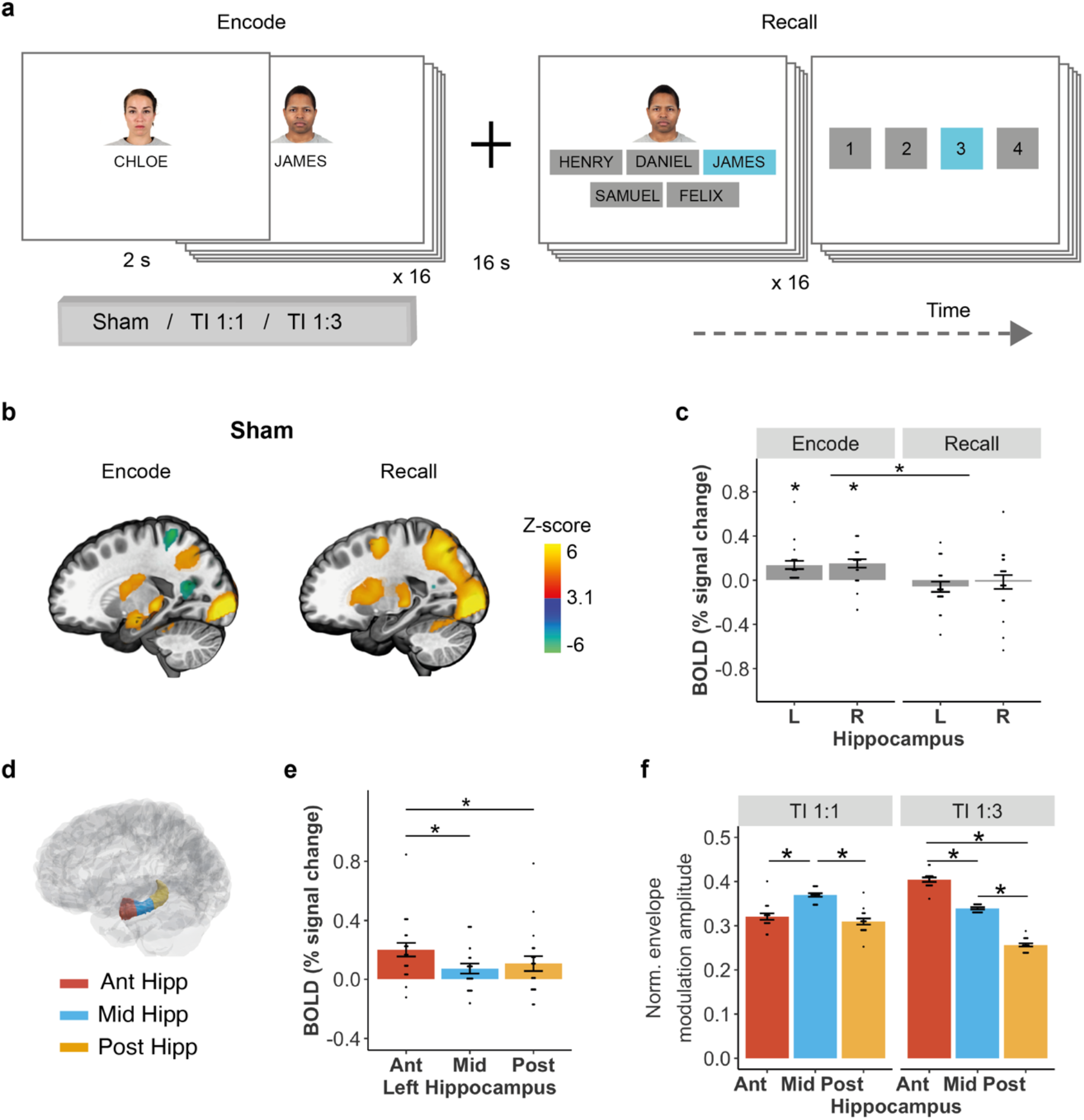
Experimental design, BOLD signal during sham and hippocampal fields. **a**, Experimental Design for Hippocampal Dependent Face-Name Memory Task. The task was composed of 9 blocks of encoding and recall. Each block contained 16 unique face-name pairs followed by a delay and a recall period, where participants tried to select the correct name of each face out of five options (i.e., one target name, two foil names that were present in the block but associated with a different face, and two distracting names that were not present during the task). After each name selection, participants were asked to rate their choice confidence (1, not confident at all to 4, extremely confident). **b**, Whole-brain group z-score change in BOLD signal during encode and recall stages of the task, showing a BOLD signal increase in the left hippocampus during the encoding, but not recall. **c**, Group median change in BOLD signal in the left (L) and right (R) hippocampi during encoding and recall stages in sham condition blocks. Showing significant BOLD signal increase during the encode, but not recall stage; see **Table S2** for full statistics. **d**, Schematic of the Ant, Mid and Post ROIs along the hippocampal longitudinal axis. **e**, Group median change in BOLD signal in the anterior (Ant), middle (Mid) and posterior (Post) regions of the left hippocampus during the encoding stage in the sham condition. Showing a larger BOLD signal increase in the Ant hippocampal region in relation to the Mid and Post regions; see **Table S3** for full statistics. **f**, Participants’ envelope modulation amplitude in hippocampal ROIs during TI 1:1 and TI 1:3 stimulations, computed with individualised MRI-based anatomical models; N=16 subjects. Showing a steering of the envelope amplitude peak from Mid hippocampal ROI during TI 1:1 stimulation to Ant hippocampal ROI during TI 1:3 stimulation; ROI amplitudes were normalised to total hippocampal exposure, see **Table S5** for full statistics. Asterisks identify significant differences, p<0.05. Bar plots show mean and standard error (SE), black dots show individual participant data. Images in **b** were thresholded at Z > 3.1, with a cluster significance level of p<0.05, and are displayed in x = -21 plane of the MNI template. N=20 throughout except for **f** where N=16. Faces in **a** were retrieved from the Chicago Face Database^68^ and reproduced here with permission from the authors under a Creative Commons Attribution 4.0 (CC BY 4.0) license.

We first examined the BOLD signal during the task without stimulation (i.e., sham condition, **Fig 2b**). As expected, the task elicited evoked BOLD activity in both hippocampi during encoding (one-sample t-test, one-sided; left hippocampus t_(19)_=3.69, p=0.002; right hippocampus t_(19)_=3.92, p=0.002; FDR corrected), but not during recall (t_(19)_=-1.28, p=0.89; right hippocampus t_(19)_=-0.25, p=0.79; FDR corrected, **Fig. 2c)**, similar to previous reports^23,27^. The LMM confirmed the significant effect of task stage (F_(1,57)_=20.492, p=3.09×10^−5^) and lack of hemisphere or interaction between the two (Fs<0.6, ps>0.4, **Table S2**). Along the hippocampal longitudinal axis (**Fig. 2d**), the BOLD signal increase in the left hippocampus during encoding was largest in the Ant region (main effect of ROI, F_(2,38)_=8.72, p<0.001; Ant– Mid, p<0.001; Ant–Post, p=0.014; Mid–Post, p=0.55, **Fig. 2e** and **Table S3**). Across the left hippocampal regions, the BOLD signal was larger when the memory association was encoded correctly (main effect of response type (correct vs incorrect), F_(1,95)_= 11.09, p=0.001 and ROI, F_(2,95)_=4.58, p=0.012, but no interaction, F_(2,95)_=0.94, p=0.39, **Table S4**). These results show left hippocampal activity is modulated during formation of correct associations, consistent with previous studies^22^. In contrast, the BOLD signal increase in the right hippocampus during encoding was similar across hippocampal regions (F_(2,38)_=0.20, p=0.82), and we did not observe a difference in BOLD signal between correct and incorrect encodings (Fs<1.4, ps>0.3, **Table S4**).

To assess whether tuning the current ratio from 1:1 to 1:3 steered the TI stimulation locus towards the Ant hippocampal region, we performed individualised simulations based on participants’ anatomical models and electrode locations (four subjects had to be excluded from the modelling since their electrodes were not visible in the MRI). We found that TI with 1:1 current ratio resulted in a relatively larger envelope modulation amplitude in the Mid region of the hippocampus (Ant: 0.32∓0.03 median∓SD relative to total hippocampal exposure; Mid: 0.37±0.02; Post: 0.31∓0.03; linear mixed-effects model (LMM) F_(2,30)_=26.05, p=2.7×10^−7^; Mid-Post/Ant, p<0.0001, Post-Ant, p=0.42, **Fig. 2f**). Changing the current ratio to 1:3 indeed steered the location with the largest amplitude to the Ant region of the hippocampus (Ant: 0.40±0.02; Mid: 0.34∓0.01; Post: 0.26∓0.02; LMM: F_(2,30)_=359.62, p<2.2 ×10^−16^; p<0.0001 across all regions, **Fig. 2f** and **Table S5**), as predicted by our simulations in the MIDA model.

### Targeted modulation of memory-evoked hippocampal activity

We next assessed whether TI stimulation affected the BOLD signal evoked in the left, i.e., targeted, hippocampus. We found that TI 1:1 stimulation did not significantly change the BOLD signal evoked by the associative memory task (**Fig. 3a** top panel and **Fig. 3b**). In contrast, TI 1:3 stimulation that was steered to the Ant region, reduced the BOLD signal (effect of stimulation F_(2,95)_=3.2, p=0.04; task stage F_(1,95)_=44.84, p=1.49×10^−9^; interaction F_(2,95)_= 2.96, p=0.056; Encode: Sham–TI 1:1, p=0.953; Sham–TI 1:3, p=0.006; TI 1:1–TI 1:3, p=0.015, **Fig. 3a** bottom panel, **Fig. 3b** and **Table S6**). The reduction in the evoked BOLD signal by the TI 1:3 stimulation in the left hippocampus was significant across hippocampal regions, with higher magnitude in the Ant region exposed to the largest relative envelope modulation amplitude, (**Fig. 3c**, effect of stimulation F_(2,152)_=12.65, p=8.13×10^−6^ and ROI, F_(2,152)_=6.35, p=0.002; but no interaction between the two p=0.76; Mean Difference: Ant_(Sham)_-Ant_(TI 1:3)_ =0.18, Mid_(Sham)_-Mid_(TI 1:3)_=0.09, Post_(Sham)_-Post_(TI 1:3)_=0.14, **Table S7**). This pattern was confirmed by inspecting the group-level contrasts comparing stimulation conditions in the left hippocampus (**Fig. 3d**). There were no voxels with significant BOLD differences between sham and TI 1:1 stimulation, whereas during TI 1:3 stimulation there was a reduction in BOLD activity predominantly in the Ant and Post segments of the hippocampus in relation to both sham and TI 1:1 stimulation (percentage voxels with significant signal change, Sham>TI 1:3: Ant=30%, Mid=2%, Post=25%; TI 1:1>TI 1:3: Ant=31%, Mid=3%, Post=25%). The amplitude of the evoked BOLD signal in the left hippocampus during TI 1:3 stimulation, but not TI 1:1 stimulation, was larger when the memory associations were encoded as in the sham condition correctly (**Fig. 3e, Fig. S3** and **Table S8**). In addition, the spatial pattern of activity in correct compared to incorrect trials was similar between sham and TI 1:3 conditions (**Fig. 3f**). This indicates that despite a reduction in BOLD signal during the TI 1:3 stimulation, the relative signal difference between correct and incorrect encodings is maintained.

**Fig. 3:**
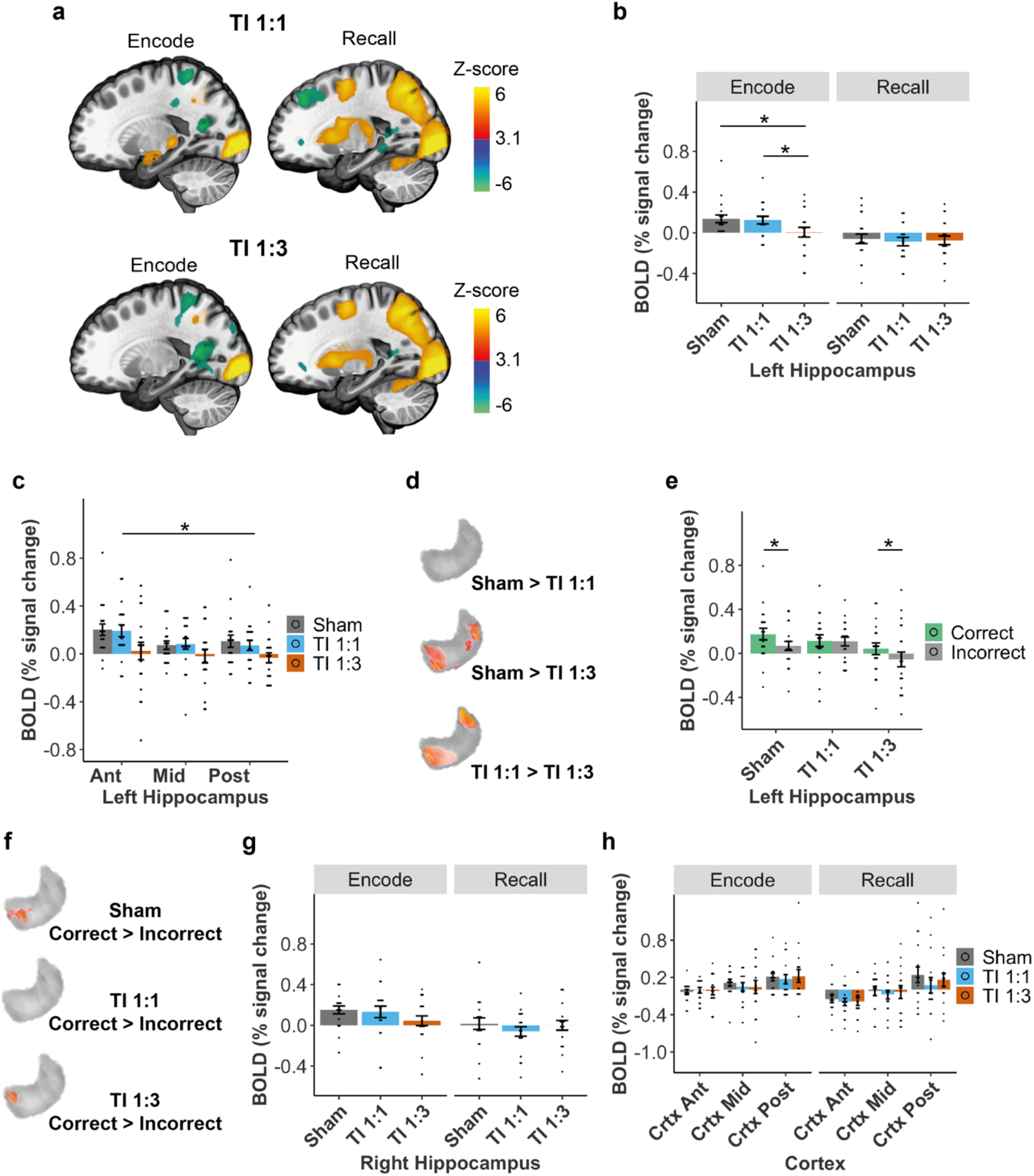
Effect of TI stimulation on hippocampal episodic memory activity. **a**, Whole-brain group z-score change in BOLD signal during encode and recall stages of the task for the TI 1:1 and TI 1:3 stimulation conditions. Note the increase in BOLD signal in the left hippocampus during encode for the TI 1:1 condition, similar to sham (**Fig. 2a**), but not for the TI 1:3 condition. **b**, Comparison of group median change in BOLD signal between stimulation conditions, in the left (i.e. stimulated hippocampus) during encoding and recall stages. Showing an effect of stimulation and reduction in the evoked BOLD signal in the left hippocampus during encoding stage by TI 1:3 stimulation; see **Table S6** for full statistics. **c**, Comparison of group median change in BOLD signal between stimulation conditions, in the Ant, Mid, and Post regions of the left hippocampus during the encoding stage; see **Fig. 2d** for ROIs schematic. Asterisk indicates a reduction in the evoked BOLD signal during the TI 1:3 stimulation across regions; see **Table S8** for full statistics. **d**, Voxelwise group-level contrasts comparing stimulation conditions in the left hippocampus, confirming higher BOLD signal for the sham and TI 1:1 conditions compared to the TI 1:3 condition. No significant voxels observed for the comparison between sham and TI 1:1, confirming the results observed in the hippocampal ROIs. **e**, Group median change in BOLD signal in the left hippocampus for memory associations encoded correctly (green) and incorrectly (grey). Showing significantly higher BOLD signal for correct compared to incorrect associations in the left hippocampus during sham and TI 1:3, see **Fig. S3** and **Table S8** for full statistics. **f**, Voxelwise group-level contrasts comparing correct and incorrect encoded associations in the left hippocampus, showing that BOLD signal during the formation of correct associations is predominantly modulated in a cluster located in the anterior portion of the hippocampus for sham and TI 1:3 conditions, while no differences between BOLD signal for correct and incorrect associations were observed for the TI 1:1 condition. **g**, Same as (**b**) but for the right hippocampus, where there is no effect of stimulation; see **Table S6** for full statistics. **h**, Comparison of group median percentage change in BOLD signal between stimulation conditions, in the anterior (Ant), Middle (Mid), and posterior (Post) regions of the overlying cortex; see **Fig. 1d** for ROIs schematic; see **Table S9** for full statistics. Showing no difference in the BOLD signal change between stimulation conditions. Asterisks identify significant differences, p<0.05. Bar plots show mean and standard error (SE), black dots show individual participant data. Images in **a** were thresholded at Z > 3.1, with a cluster significance level of p<0.05, and are displayed in x = -21 plane of the MNI template. Images in **d** and **f** were thresholded at significance level of p<0.05, voxelwise permutation-based t-tests on the ROI. N=20 throughout except for **h** where N=16.

Next, we assessed whether the BOLD signal was modulated in the right (not targeted) hippocampus and the overlaying cortical regions underneath and between the stimulation electrodes. There were no significant differences in BOLD signal in the right hippocampus (**Fig. 2g**, Fs<2, ps>0.1, **Table S6**) or left ROIs closed to the stimulation electrodes (**Fig. 2h**, ps>0.7, **Table S9**). To support that the lack of changes in BOLD signal were not driven by anatomical variability (electrode locations determined using landmarks based on head size) or reduced sample (16 out of 20), we proceeded to extract BOLD signal in the left temporal lobe (excluding the hippocampus) for all participants. There was no significant effect of stimulation on BOLD signal in the left temporal ROI (ps>0.2, **Table S9**), which was again confirmed by group-level voxelwise analysis. As a final interrogation on the spatial specificity of the BOLD changes, we investigated whether BOLD signal was modulated by stimulation in the left amygdala, located anteriorly to the hippocampus. We did not observe changes in BOLD signal for the left amygdala (**Fig. S4**). The lack of BOLD signal change in the overlaying cortical regions and neighbouring amygdala cannot be explained by a lack of task activation since, across conditions, we observed evoked BOLD signal in these regions (**Table S13**).

Finally, we tested whether the change in BOLD signal was mediated by a difference in the participants’ memory performance – earlier studies have shown that stimulus-induced BOLD is modulated by background activity^28,29^ and behaviour differences can mediate BOLD signal differences^30^. We found that the participants recall accuracy was above chance for all stimulation conditions (ts>10, ps<0.001, **Table S11**), but there was no difference in the recall accuracy or recall time between stimulation conditions (**Fig. S5** and **Table S12**). There was a small difference in the confidence rating, explained by a general increase in confidence during the TI 1:3 condition, but no interaction between stimulation condition and response type.

Taken together, our results demonstrate a non-invasive focal modulation of evoked neural activity in the targeted hippocampus. The hippocampal decrease in BOLD signal observed during the TI 1:3 condition is in alignment with previous animal studies, showing theta frequency stimulation decreases the magnitude of the BOLD signal in the hippocampus^31-33^. This would suggest that larger field magnitudes should result in larger decreases in BOLD signal. We observed some evidence to support this relationship (significant Pearson, but no significant robust correlations possibly due to the small sample size) in the same hippocampal regions where BOLD signal was mostly modulated by the stimulation (‘Ant’ and ‘Post’ regions, **Fig. S6**).

### Modulation of hippocampal functional connectivity

Given that successful associative memory involves interactions between the hippocampus and cortical networks, in particular the antero-temporal (AT; more connected to the Ant hippocampus) and posterior-medial (PM; more connected to the Post hippocampus) networks^34,35^, (**Fig. 4a**), we sought to explore whether stimulation of the hippocampus changes the functional connectivity (FC) in those networks. In the absence of stimulation, successful encodings increased FC between the Ant and Mid, but not Post, hippocampal regions and the AT network, (**Fig. 4b**, AT: Ant: t=2.322, p=0.022, uncorrected; Mid: t=3.117, p=0.029, FDR-corrected, **Table S13**). There was no change in FC between hippocampal regions and the PM network or during recall.

**Fig. 4:**
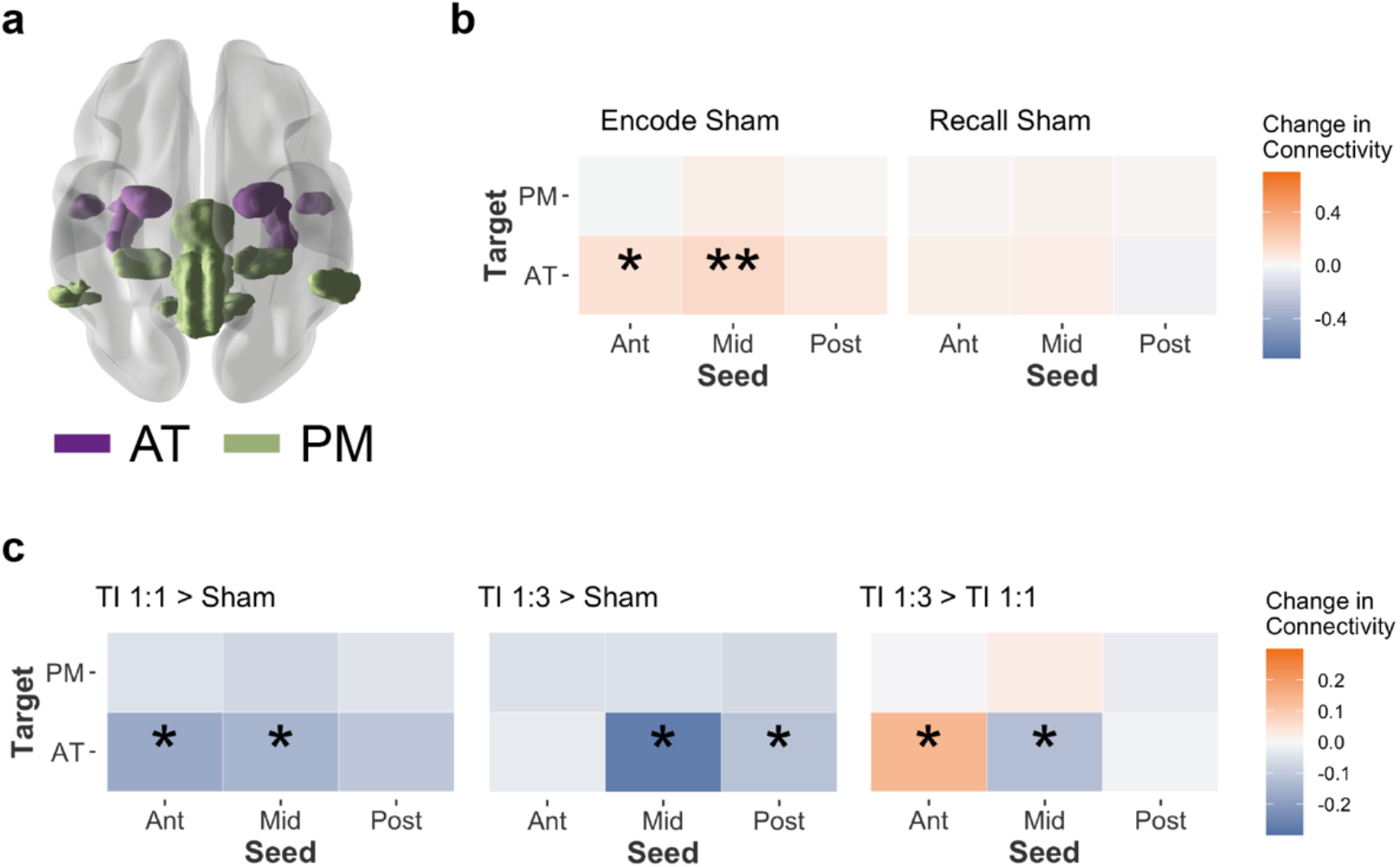
TI stimulation change in hippocampal-cortical functional connectivity. **a**, Antero-temporal (AT, purple) and posterior-medial (PM, green) hippocampal-cortical networks. **b**, Group mean change (% signal change) in functional connectivity between the anterior (Ant), Middle (Mid), and posterior (Post) regions of the left (L) hippocampus and the AT and PT networks during encoding and recall task stages in sham blocks (i.e., without stimulation) for the contrast correct > incorrect. Showing a larger connectivity between the Ant and Mid regions of the L hippocampus and the AT network during encoding of successful associations; one-sample t-tests; N=20; **, p < 0.05 FDR-corrected; *, p < 0.05 uncorrected; see **Table S13** for full statistics. **c**, Effect of TI stimulation on functional connectivity, showing the same as **b** but comparing changes between stimulation conditions during encoding stage in which a connectivity increase was observed in sham blocks. Showing a reduction in functional connectivity by both TI 1:1 and TI 1:3 stimulations (relative to sham), and when comparing the TI stimulations, a larger connectivity in the Mid region during TI 1:1 and in the Ant region during TI 1:3 (LMM with significant 3-way interaction between stimulation type, seed and network; *, p < 0.05 corrected post-hoc contrasts; see **Table S13** for full statistics).

Compared to sham, both TI stimulations reduced FC between the hippocampus and the AT network. This reduction was localised to the Ant and Mid regions of the hippocampus during TI 1:1 stimulation and the Mid and Post regions during TI 1:3 stimulation (**Fig. 4c**, significant interaction between stimulation type, seed and network F_(4,1576)_=2.5, p=0.04; post-hoc tests, TI 1:1: Ant, p=0.8; Mid, p<0.001; Post, p=0.04; TI 1:3: Ant, p=0.001; Mid, p=0.006; Post, p=0.07). Comparison of FC between TI stimulation conditions showed a higher relative connectivity at the hippocampal region that was exposed to the largest envelope modulation amplitude. Specifically, FC between the Mid hippocampus and the AT network was larger during TI 1:1 stimulation than during TI 1:3 stimulation (p=0.023, **Fig. 4c**). In contrast, FC between the Ant hippocampus and the AT network was larger during TI 1:3 stimulation (p=0.008, **Fig. 4c** and **Table S13**). We did not find specific seed-network FC differences between the stimulation conditions during the recall period; however, we found a main effect of stimulation, explained by lower connectivity values during the TI 1:1 stimulation (F_(2,1576)_=8.320, p=2.544×10^−4^, **Table S14**).

These results suggest that the reduction in the memory evoked BOLD signal in the hippocampus occurred alongside a reduction in the functional connectivity between the hippocampus and its AT cortical network. The concurrent reduction in the hippocampal BOLD signal and its functional connectivity may indicate a lower metabolic cost of the underlying memory operation^36^.

### Enhancing hippocampal-dependent episodic memory performance

We next aimed to explore whether the TI stimulation of the hippocampus could affect the underlying memory function. We tested a new cohort of twenty-one participants (mean age 23.2 ± 3.7 SD, 10 females) with a similar hippocampal-dependent face-name paired associative task but this time with a more extended period of stimulation and a larger number of behaviour trials (**Fig. 2a**) – an experimental protocol designed to probe behavioural effects of stimulation^37^. We applied the TI 1:3 stimulation that showed stronger modulation of hippocampal memory BOLD signal and sham in counterbalanced order in two separate experimental sessions. In contrast to the first experiment, we applied the stimulation continuously not just during encoding but also during the maintenance and recall periods. In addition, since earlier studies have shown that retrieving a memory can transform the information being remembered^38,39^ thereby facilitating or impeding the memory^40^, we explored this effect by retesting all the face-name pairs again after 30 min.

We found an effect of TI stimulation on participants’ performance (GLMM: 𝒳^2^(2)=6.353, p=0.042; **Fig. 5a**). Specifically, participants showed higher proportions of correct (i.e., target) recalls during TI compared to sham (p=0.007), with no difference in the number of foils (p=0.142) or distractors (p=0.384). We confirmed the higher recall accuracy during TI stimulation employing a frequentist binomial model of correct vs incorrect responses (𝒳^2^(2)=5.857, p=0.016), and Bayesian model, showing posterior distribution estimation of 12% mean accuracy improvement (mean estimate =0.12, 95% credible interval (CI) =0.02 to 0.21; 99.15% of the posterior >0, **Table S15**). The total stimulation duration differed slightly between participants (mean 34.5 ±3 SD min) due to self-paced responses. However, recall accuracy was not correlated with stimulation duration (r=-0.045, p=0.85; Pearson correlation). Comparison of accuracy at recall and re-test, for items that were correctly remembered at recall, showed an effect of stimulation (𝒳^2^(1)=7.581, p=0.006) and an effect of time of testing (𝒳^2^(1)=233.124, p<0.001), but no interaction (𝒳^2^(1)=0.063, p=0.802), indicating that the memory benefit gained during TI stimulation was maintained at re-test (**Fig. 5b**).

**Fig. 5:**
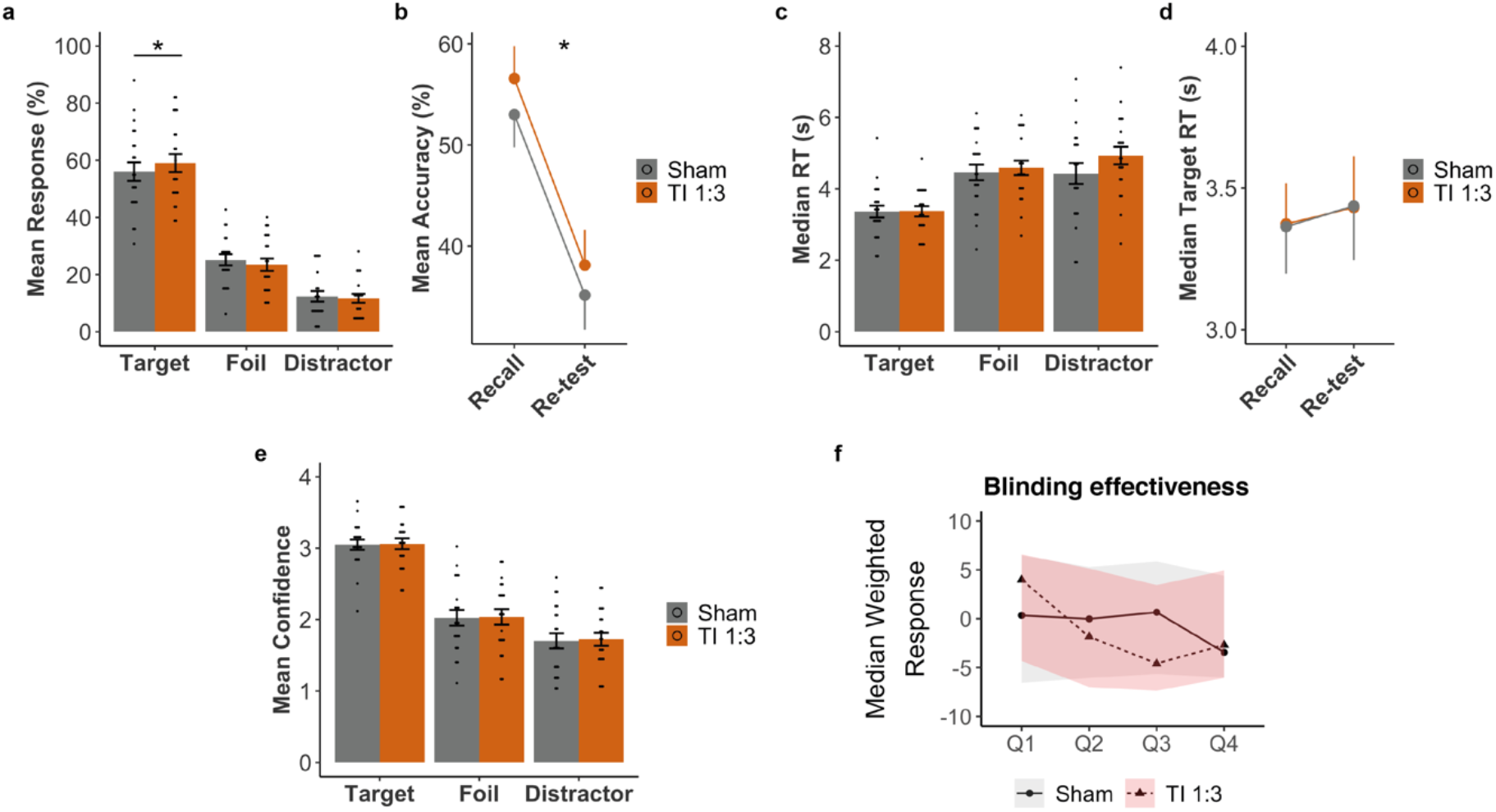
Probing the effect of hippocampal TI stimulation on behavioural function. **a**, Comparison of participants’ memory performances during recall between sham (grey) and TI 1:3 (orange) across response type (target, foil and distractor), showing higher probability of target responses, i.e. face-name pairs correctly remembered. **b**, Comparison of mean accuracy for target selections between recall and re-test (30 min after first recall) for sham (grey) and TI 1:3 (orange), showing that target selection was higher for TI 1:3 condition at both time points. **c**, Same as **a** but for median reaction time, showing no differences between stimulation conditions. **d**, Same as **b** but for median reaction time for target responses, showing no significant difference between recall and re-test or stimulation conditions. **e**, Same as **c** but for mean confidence ratings, which were similar between stimulation conditions. **f**, Blinding effectiveness. Shown are median weighted scores and 95% confidence intervals for TI 1:3 (pink) and sham (grey) conditions; positive score, participants reported having stimulation, negative score not having stimulation. Confidence scores between 1 (not confident) and 10 (extremely confident). Confidence intervals overlapped throughout indication that participants were blinded to the stimulation condition. **a-e**, Bar or dot plots show mean and standard error (SE), black dots show individual participant data. Asterisks (*) indicate significant differences between stimulation conditions. See **Table S15** for full statistics. N=21 throughout.

The improvement in recall accuracy was not accompanied with a change in recall time, (**Fig. 5c**, GLMM/LMM: no main effect of stimulation using multinomial logistic regression 𝒳^2^(1)=3.017, p=0.082, or binomial model 𝒳^2^(1)=2.993, p=0.084; no interaction between stimulation and recall time, ps>0.2). Comparison between median reaction times for recall and re-test showed no effect of stimulation, time or interaction (**Fig. 5d**, 𝒳^2^s<0.6, ps>0.4). Similarly, there was no effect of stimulation on confidence rating during recall (**Fig. 5e**, no main effect of stimulation 𝒳^2^(1)=0.35, p=0.557, or interaction between confidence per response category and stimulation p=0.809, **Table S15**).

### Safety, tolerability, and blindness

Across both experiments, participants tolerated well the TI stimulation. There were no adverse effects and only a few incidences of mild common side effects (see **Table S16** for fMRI experiment and **Table S17** for behavioural experiment). In the last experiment, because stimulation and sham were performed in separate sessions, we could directly compare the incidence of the side effects. We observed that only itchiness at the electrode site was higher for TI than sham stimulation (Z= -2.354, p=0.019, **Table S17**). Despite the high current densities the threshold current intensity at which participants reported perceiving extraneous skin sensation underneath the electrodes was much higher than when we tested short conventional transcranial alternating current stimulation (tACS) during setup (mean ± SD, tACS: 0.424±0.195 mA; TI: 2±0.540 mA; t_(65)_=17.203, p<0.001, unpaired two-tailed t-test, N=42 reporting perceptual sensations during tACS and N=25 during TI, pooled from both studies, see **Tables S18-S20**).

Finally, to assess stimulation blindness, we asked participants to report whether they felt they were stimulated (‘yes’/’no’ response) and their answer’s confidence, as in^41^, at four time points during each session of the behavioural experiment. We found that participants were blinded to the stimulation condition, i.e., no difference between TI stimulation and sham in participants’ weighted confidence of receiving stimulation (**Fig. 5f**).

## Discussion

In this paper we present the first demonstration of non-invasive electrical DBS in humans using TI of kHz electric fields, expanding on our earlier validation in rodents^14,42^. We first use electric field modelling and measurements in a human cadaver to verify that the locus of the transcranial TI stimulation can be steerably localised to the human hippocampus with minimal exposure of the overlying cortex. We then use neuroimaging and behaviour experiments in healthy humans to demonstrate focal non-invasive modulation of hippocampal memory activity and the capacity to augment memory performance.

Our modelling predicts that the TI fields locus in the hippocampus had a median amplitude of ∼0.25 V/m per ∼0.45 mA/cm^2^ applied current density (1 mA current in our electrodes), consistent with previous computational studies^43-46^. Yielding a median hippocampal envelope amplitude of ∼0.5 V/m in our human studies (∼0.9 mA/cm^2^ applied current density) and ∼0.2 V/m difference between the Ant region and the Mid/Post regions. Similar field amplitudes have been consistently reported to synchronise neural spiking activity tuned to the endogenous oscillation frequency range in-vitro^47^ and in-vivo^48-50^. One limitation of our cadaver measurements is that data was collected at a temperature lower than the living body, resulting in lower tissue conductivity and, consequently, higher electric field amplitudes for fixed current densities^50^. However, since the electric field amplitudes change equally across the head tissues^50^, the relative field distribution estimations in the cadaver were not affected.

Our neuroimaging experiments aimed to probe a focal physiological effect. We demonstrate that when the TI stimulation locus is transiently applied to the hippocampus with a theta-band difference frequency during encoding of episodic memory, it reduces the hippocampal evoked BOLD signal without affecting evoked BOLD signal in the overlying cortex. The BOLD reduction was strongest when the TI stimulation locus was steered to the anterior hippocampus, in line with the repeated reports on the central role of this region in the successful encoding of face-name associations^20,22,23^. The evidence of focal physiological effect in the hippocampus without behavioural memory differences, enables robust inference of target engagement since memory differences are associated with widespread changes in BOLD signal^51^ and can be mediated by remote stimulation of functionally connected sites^52^ or by stimulation-independent changes in brain state^53^.

Our subsequent experiment aimed to probe the behaviour consequence of the physiological effect in the hippocampus. We demonstrate that when the TI stimulation is applied to the hippocampus for a more extended period throughout the episodic memory operation, it provides an improvement in memory accuracy. The magnitude of the memory improvement was in line with earlier TES studies modulating working memory performance^37^. We observed that memories formed during TI endured the effects of re-test, but the rate at which they were forgotten was similar to sham. Future studies aimed at understanding longer-term effects of stimulation in memory should investigate whether more extended continuous TI stimulation and/or repeated sessions, may be able to achieve stronger and sustained memory benefits. Those studies may be able to pinpoint the optimal stimulation timing, i.e., memory encoding, maintenance, and/or recall.

What is the possible mechanism by which the theta-band TI stimulation of the hippocampus reduces its memory evoked BOLD response and augments its function? A strong body of evidence shows modulations in theta oscillations in the medial temporal lobe during episodic memory^54^. Since TI stimulation modulates neural activity at the difference frequency of the kHz-frequency electric fields^14^, we anticipate that its application with a theta-band difference frequency will modulate endogenous theta oscillations in the local hippocampal network, resulting in a change in BOLD signal. Whilst the direction of neuronal activity cannot be unambiguously inferred from BOLD responses, and discrepancies in the relationship between BOLD signal and neural activity across cortical and hippocampal regions are well documented^55^, our observed reduction of hippocampal BOLD signal due to the TI stimulation is consistent with previous reports in animal models delivering theta frequency electrical stimulation to the hippocampal circuit^31-33^. Multimodal data demonstrated that a reduction in hippocampal BOLD signal can be observed alongside increased blood volume and increased spiking activity as a consequence of local increases in deoxyhaemoglobin^32,56,57^. Although our current data does not allow us to disambiguate which components of the BOLD signal are affected by TI stimulation, it is unlikely that the effect observed can be simply explained by neurovascular changes. This is because no changes in BOLD signal were observed when TI was directed to the Mid hippocampus. Further, changes in functional connectivity were specific to the AT cortical network and their modulation between the two TI conditions was in line with TI fields’ distribution along the hippocampal longitudinal axis, providing further evidence of the TI stimulation specificity and steerability. Studies using intracranial electroencephalographic (iEEG) recordings from the hippocampus will be able to elucidate the direct neural response and offer further mechanistic insights to the observed BOLD changes.

Could the change in hippocampal BOLD signal and memory performance have been mediated by stimulation of the overlying cortex exposed to larger field amplitudes? The lack of stimulation effect on the evoked BOLD signal in the overlying cortex and the fact that the participants’ change in hippocampal BOLD signal was correlated, albeit weakly, with the envelope modulation amplitude in the hippocampus but not in the overlying cortex render this possibility unlikely. Could the observed change in hippocampal BOLD signal and memory performance have been mediated by a conventional AC stimulation of the kHz fields in the hippocampus? A recent electrophysiological study investigating the effect of kHz-frequency electric fields in hippocampal brain slices reported no effect on neural activity, even with amplitudes that are two orders of magnitude larger than those we used in this study^58^, consistent with our earlier electrophysiological study in live mouse brain^14^.

Overall, TI stimulation was well tolerated, no adverse effects were recorded, and side effects were mild. We used current densities that are considered safe and in line with those applied across non-invasive electrical stimulation studies, where no serious adverse effects have been reported^59^. We found that the thresholds at which TI stimulation produces extraneous sensations are much higher than those for conventional tACS and allowed for adequate stimulation blinding. This is useful as therapeutical applications might benefit from higher current densities for which blinding becomes harder to achieve^60^.

The hippocampus is important in a myriad of brain functions, including learning and memory, spatial navigation, and emotional behaviour. It also plays a central role in many of the most common brain disorders, including Alzheimer’s disease, epilepsy, depression and schizophrenia^61^. By modulating hippocampal neural activity noninvasively, TI stimulation offers new opportunities to probe its causal role in brain functions. Future studies using different electrode configurations and current parameters may sculpt the TI stimulation locus to focally modulate the neural activity in other deep brain structures. Tuning the difference frequency of the applied kHz-frequency fields will allow exploring the frequency band specificity of the target brain regions and contribute to inform stimulation optimisation strategies to treat brain disorders.

## MATERIALS AND METHODS

### Electric field simulations

To characterize the in-vivo exposure to high frequency fields and to low-frequency TI modulation, as well as for the identification of optimized stimulation configurations, dosimetric electromagnetic simulations involving anatomical head models were performed. Two kinds of head models were used: 1) a highly detailed and accurate reference head model, and 2) personalized head models that permit to study the relationship between inter-subject anatomy and exposure variability and subject-specific BOLD response.

### Reference Head Model

For maximal simulation realism, the highly detailed MIDA head model (jointly developed with the FDA^18^ was used. This model is based on high-resolution (< 0.5 mm throughout) multi-model MRI data, which allows for the distinction of more than 100 different tissue regions, including a range of deep brain targets, the crucial distinction of cancellous and cortical skull layers, the dura, various scalp layers (skin, muscle, fat, tunica) and the complex distribution of cerebrospinal fluid, among other tissues. Co-registered DTI data provides the necessary information about brain heterogeneity and anisotropy, as well as the local principal orientation of fibres.

### Personalised Head Models

Individualised (though less accurate and detailed) head models were generated from T1 images (see MRI Data Acquisition) using the SimNIBS framework (version 3.2^62^), employing the ‘headreco’ pipeline^63^ to distinguish six tissue classes: scalp, skull, cerebrospinal fluid, grey matter, white matter, and head cavities. Segmented images were visually inspected and manually corrected when necessary (manual corrections were mostly restricted to the skull-CSF boundary). Because the hippocampi are not included in the automatic segmentation from SimNIBS, these were extracted using FreeSurfer (see Regions of Interest) and converted into tissue surfaces using the iSEG software (IT’IS Foundation, Zurich, Switzerland). Using subject-specific images also permitted to accurately position the scalp electrodes on the reconstructed scalp surfaces in 16 out of 20 participants from the fMRI experiment (in 4 participants the field-of-view of the T1 image did not allow for a clear localization of the electrodes on the skin).

### Electromagnetic field computation

The reference and the reconstructed subject-specific head models were imported as surface mesh entities into the Sim4Life (ZMT ZurichMedTech AG, Zurich, Switzerland) platform with extended TI modelling functionality. Electrode geometries (2 cm diameter cylinders) were created in Sim4Life, placed at the identified electrode locations, and aligned to the head surfaces, while ensuring gap-less contact. The setup for EM simulations consisted of dielectric property and boundary condition assignment, followed by gridding and voxeling for numerical discretization. The simulations were executed using Sim4Life’s finite element method (FEM) low frequency electro-quasistatic, ohmic-current-dominated (EQSCD) solver, that computes solutions to the quasistatic and ohmic-current-dominated approximation of the Maxwell equation (∇σ∇φ, with boundary conditions) on an adaptive, rectilinear grid, where φ is the electric potential and σ the electric tissue conductivity distribution. EQSCD assumes that ohmic (resistive) currents dominate over displacement currents at the frequencies of interest and that the wave-length is large compared to the computational domain^64^ – conditions that were confirmed by a solver-performed analysis. The conductivities of the non-brain tissues were assigned based on a recent meta-analysis of reported human head electrical conductivity values^65^. To account for the important impact of brain tissue dielectric anisotropy and heterogeneity, DTI-based electrical conductivity tensor maps were generated. The local main orientation was derived through spectral decomposition of the DTI tensors and in turn combined with the longitudinal and transversal conductivities according to^66^, to reconstruct σ tensors. A convergence analysis was performed to identify an optimal grid resolution that ensures sufficient accuracy (i.e., negligible discretization errors) while minimizing the number of discretization elements (voxels) to reduce computational resource requirements. Simulations were executed at a homogeneous 0.5 mm resolution, which resulted in models consisting of about 80 million voxels. Each TI stimulation exposure condition required the execution of two EM simulations per subject, one for each electrode pair. Dirichlet boundary conditions were assigned to the active electrodes (thus capturing the inhomogeneous current distribution across the electrode interface), applying an arbitrary voltage difference of 1V subject to subsequent current normalization.

### Computed TI exposure metrics

The calculated electric (E) fields for each electrode pair were normalized to an input current of 1 mA, by integrating the normal current density over a surface surrounding an electrode. The spatial distribution of the projected TI envelope modulation amplitude along the local brain structure orientation 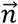 was computed from the fields of both electrode pairs using 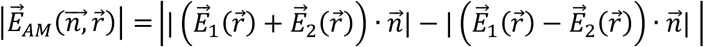, where 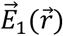and 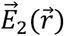 are the fields generated by the first and second electrode pair, respectively, at the location 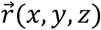. The local brain structure (e.g., white-matter fibres, organized pyramidal neurons in the hippocampus) orientation was identified as the principal axis of the corresponding DTI tensor.

### Electric field measurements in human cadaver

A human male cadaver (93 years old) with no known brain disorder was provided by the “service des corps donnés à la science” by Aix Marseille Université. Experiments were performed in the Faculty of Medicine La Timone (Aix Marseille Université). The subject was perfused with zinc chloride, stored in freezer until the experiments and left to warm to 20°c before the recording session.

Electric fields were recorded using three stereoelectroencephalography (sEEG) 15-contact electrodes, ring diameter 0.8 mm, length 2 mm, useful exploration length 51 mm (Alcis, Besançon, France, 2069-EPC-15C-35). The sEEG electrodes were implanted in the left mesial temporal lobe, perpendicular to the hippocampal longitudinal axis. The technique of implantation was based on the neurosurgeon’s experience in performing SEEG (RC) and was similar to the one routinely applied to human patients for the presurgical work-up of drug-resistant epilepsy. Each electrode was orthogonally inserted through a short 20-mm guidance screw (Alcis, Besançon, France, 2023-TO-C-020) after 2.5 mm diameter drilling of the bone. Reference and ground electrodes were placed on the shoulder of the cadaver using ECG electrodes (WhiteSensor WS, Ambu® Inc., MD, USA, 1.5×1.5 cm).

The electric potential signals from the sEEG electrodes were amplified and sampled at 30 kS/s using the RHS Stim/Recording Controller (Intan Technologies, Los Angeles, CA, USA). The stimulating currents were applied using 1.5 cm x 1.5 cm scalp electrodes (WhiteSensor WS, Ambu® Inc., MD, USA) as in **Fig. 1b**. The two currents were generated using two electrically isolated current sources (DS5, Digitimer Ltd., UK) driven by voltage waveforms from a function generator (Keysight, EDU33212A, Santa Rosa, California, USA). In the case of TI stimulation, we applied one current at 2.005 kHz frequency and 1 mA (current density ∼0.45 mA/cm^2^) amplitude and a second current at 2 kHz frequency and same amplitude (i.e., TI with Δf = 5 Hz and 1:1 current ratio, **Fig. 1c**, left). In the case of conventional tACS, we applied two currents at 5 Hz frequency and 1 mA amplitude. Each stimulation condition was applied for 25 s. The 3D location of the electrodes within the expected mesial temporal anatomical targets was confirmed by a CT of the head at the end of the experiment. The recorded data were analysed using a custom-written script in MATLAB (Mathworks, Natick, MA). The electric potential signals were first bandpass filtered using a 1st order Butterworth filter with cut-off frequencies of 0.5 kHz and 5 kHz in the case of TI stimulation and 1 Hz and 40 Hz in the case of conventional tACS. The normalised envelope modulation amplitude in each electrode was estimated by first extracting the recorded signal’s envelope waveform using a Hilbert transform and a low-pass filter (i.e., 1st order Butterworth filter with a cut-off frequency of 0.5 kHz) and then computing the mean half difference between the waveform maxima and minima (averaged first in 1 s epochs and then across the 25 epochs). The envelope modulation amplitude of each electrode was then normalized to the largest envelope modulation amplitude across the electrode’s contact points. The envelope modulation ratio was estimated by dividing the amplitude of the envelope waveform by the maximum signal amplitude. The field’s envelope modulation amplitude along the axis of the recording electrodes (i.e., perpendicular to the hippocampal longitudinal axis) was estimated by computing the difference in envelope modulation amplitudes between neighbouring contact points and dividing the value by the inter contact distance. The normalised absolute amplitude in each electrode was estimated by computing the median value of the signal maxima (again averaged first in 1 s epochs and then across the 25 epochs). The field’s amplitude along the axis of the recording electrodes (i.e., perpendicular to the hippocampal longitudinal axis) was estimated by computing the difference in amplitudes between neighbouring contact points and dividing the value by the inter contact distance. The spatial maps of the normalised envelope modulation amplitude and normalised absolute amplitude were created by first applying a 3-point moving average over the electrode contacts followed by a linear interpolating of the electrode contact values of 100 × 151 grid.

Hilbert transform was applied to TI data, followed by low-pass filtering (500 Hz) using a first order zero-phase, forward-reverse Butterworth filter. Maximum and minimum amplitudes were computed by calculating the median values extracted from 1 s epochs. The amplitude envelope modulation for TI data was calculated using the average of the maximum and minimum amplitudes. The envelope modulation ratio was calculated as the ratio of the envelope modulation amplitude to the maximum amplitude. Field strengths were calculated using the first spatial derivative of the envelope modulation amplitude or maximum amplitude.

### Human subjects – in-vivo experiments

Twenty-two healthy volunteers were recruited for the MRI experiment and twenty-one for the experiment probing the effects of TI stimulation on behaviour. In the fMRI experiment, two participants were excluded, one because of technical difficulties with the MRI scanner (no images were collected) and another due to excessive movement in the scanner. Thus, the final cohort for this experiment was composed of twenty subjects (11 females, age range: 20 to 54 years old, mean age 27.1 ± 7.6 SD, 19 right-handed and 1 left-handed). For the behavioural experiment, all participants were included in the analysis (10 females, age range: 19 to 32 years old, mean age 23.2 ± 3.7 SD, all right-handed). All subjects were educated to degree level or above with no self-reported history of neurological or psychiatric illness. Participants gave written informed consent. The study conforms to the Declaration of Helsinki and ethical approval was granted by the Imperial College Research Ethics Committee (ICREC). MRI data was collected at Imperial College London clinical imaging facility (CIF).

### Face-name task

The task was designed using the Psychtoolbox^67^ for Matlab (Mathworks, Natick, MA). In the MRI experiments, responses were collected with a response box (NordicNeuroLab, Norway) that was connected to the stimulus presentation PC through a fibreoptic. In the behavioural experiment, responses were collected using a computer keyboard connected to the stimulus presentation PC. The order of the stimulation was counterbalanced across participants using a balanced Latin square. This allowed us to keep factors of no interest fixed (i.e., difficulty of a specific block or tiredness), while controlling for the variable of interest, i.e. stimulation condition.

The Face-Name task was chosen based on a strong body of evidence demonstrating that face-name associations are dependent on hippocampal function and elicit bilateral hippocampal activations in healthy subjects^20,22,23^. Faces were retrieved from the Chicago Face Database v.2.0.3^68^ and names from the Office for National Statistics (Baby Names, England and Wales, 1996; which corresponds to the dataset closest to the mean age for the faces in the Chicago Face Database, mean age = 28 years old). We selected names that had between 5 and 7 letters. Names present in both female and male lists were removed (e.g. Charlie) and if the same name was present with a different spelling (e.g. Elliot and Elliott) only the one with the highest frequency was kept. The task was composed of 9 blocks in the fMRI experiment and 12 blocks in the behavioural experiment, each containing 16 unique face-name pairs of different ethnicities (4 black female, 4 black male, 4 white female and 4 white male per block; all with neutral facial expressions). The task was composed of an encoding and a recall stage. During the encoding stage each face-name pair was displayed for 2 s. Faces were displayed in the centre of the screen with the name underneath (**Fig. 2a**). Participants were instructed to read the name underneath the faces and try to learn each face-name pair. This was followed by either a delay period (16 s) where a fixation cross was present in the centre of the screen in the fMRI experiment, or a distractor task (40 s) where participants made odd/even judgments for random integers ranging from 1 to 99 (to prevent maintenance of information in working memory) in the behavioural experiment. In the recall stage, participants were shown each face with 5 names underneath: the target name, two distractor names (i.e., names that were not present in the block), and two foil names (i.e., names that were present in the block but associated with a different face) - target and distractor names were selected to have a similar name frequency. Each name appeared in black font, inside a grey rectangle and the temporally selected rectangle was coloured in cyan. Participants moved between rectangles (using left and right index buttons in the fMRI experiment or left and right arrow keys in the behavioural experiment) and pressed a key to confirm their selection (right thumb in the fMRI or space bar in the behavioural experiment). This was followed by a confidence rating, in which participants were asked to rate how confident they were in their selection from 1 to 4 (1 being not confident at all and 4 extremely confident). Selection was made using the same procedure described for the name selection. For the recall stage, participants were instructed to respond as quickly and as accurately as possible. There was a time limit (20 s in the fMRI experiment and 8 s in the behavioural experiment) to select each name and to rate the confidence level (5 s in the fMRI experiment and 3 s in the behavioural experiment). The order of the blocks was kept constant across participants, but the order of the face-name pairs was pseudo-randomised across participants, such that no more than three faces of the same type appeared in a row. The order of face recall was randomised across participants, and the last two encoding trials were never presented at the beginning of the recall. The position of the names in the recall stage was also randomised. In the behavioural experiment, participants performed a re-test, 30 minutes after the last recall block. During this period participants were asked to recall the names matching all faces presented for that session. Stimuli were grouped in their original blocks, but blocks presented in a different order from the recall after encoding and the order of the stimuli randomised per block. The order of the blocks in the re-test was kept constant between sessions, with unique stimuli presented in each session. Confidence ratings were not collected during re-test. Participants watched a nature video between the last block of the recall period and the re-test. There was one video per session and all participants watched the same video.

### Behavioural analyses

Three main variables of interest were analysed, i.e., response type, reaction time for name selection and confidence level (reaction times for confidence were also recorded but not analysed). For each face-name trial four response types were possible at the recall stage: 1) correct association, selection of the correct name for the face presented; 2) foil (incorrect association), selection of a name that was present in the same block but did not match the face (2 foils were present per recall trial); 3) distractor (incorrect association), selection of a name that was not associated to any face across all blocks (2 distractors were present per recall trial); 4) omission, participant did not select a response within the time limit.

We first assessed accuracy across the whole task (correct vs incorrect associations) to check whether any participant had an overall performance below chance (20%), which would exclude them from further analyses. All participants were above chance (fMRI experiment: mean = 49.97%, SD = 9.77%, range 32.64 – 70.83 %; behavioural experiment – recall: mean = 58.47%, SD = 14.73%, range 33.85 – 91.15%). The distribution of responses for correct associations, foils and distractors across the whole task followed the expected pattern, with higher percentage of responses for correct associations than foils than distractors, indicating appropriate engagement with the task (fMRI experiment: correct associations = 50%, foil = 33.3%, distractor = 16.5%, omission = 0.17%; behavioural experiment – recall: correct associations = 58.46%, foil = 24.76%, distractor = 12.29%, omission = 4.49%). The number of omissions was considered negligent and removed from the dataset. We then plotted reaction times across the whole task; this showed a right-skewed distribution, typical for this metric. To trim the distribution, we calculated the 1 and 99 percentiles across all trials and participants and dropped trials below or above these thresholds.

To investigate whether the number of responses differed per response type across stimulation conditions we performed a multinominal logistic regression on the trial-by-trial data using the multinom function in the *net* package in R^69^. The outcome variable response type contained three levels, target, foil, distractor, and the level “target” was used as the intercept, with predictors for stimulation type (sham, TI 1:1, TI 1:3 in the fMRI experiment and sham and TI 1:3 in the behavioural experiment), *p* values were calculated using Wald tests. In addition, as we were interested in investigating the contrasts for correct and incorrect responses in the imaging data, we defined a binomial generalised linear mixed model (GLMM) using a logistic link function to model the effect of stimulation type on accuracy (correct vs incorrect associations). The final model included random intercepts for participant and task block (task block was not a variable of interest, as blocks were counterbalanced across stimulation conditions and was included for appropriately modelling variance in the data). In the behavioural experiment, we included in addition random intercepts for session, and modelled accuracy used Bayesian mixed effect models using the *brms* package^70^. Bayesian frameworks are robust to potential violations of normality or homoscedasticity and allow considering whether an effect is credibly different from a null value. The Bayesian model included random intercepts for participants, session, and blocks.

To investigate whether reaction times differed per response type across stimulation conditions the data was modelled with GLMM using an inverse Gaussian distribution with the identity link function^71^. The final model included random intercepts for participants and blocks in the fMRI experiment and for participants, session and trial nested in blocks in the behavioural experiment. We also run an additional model using accuracy instead of response type, to investigate whether reaction times differed by accuracy and stimulation type (again employing the inverse Gaussian distribution with the identity link function and random intercepts as described above).

Finally, we investigated the participants confidence levels for the selected face-name associations. First, we removed trials where participants did not specify a confidence level within the time limit (fMRI experiment: 0.28% trials; behavioural experiment: 0.78% trials). To investigate whether confidence levels (ordinal variable) differed per response type across stimulation conditions, the data was modelled using a cumulative link mixed model (logit link function) using the “clmm” function from the *ordinal* package in R^72^.

### Statistical procedures

All statistical analyses were conducted using R version 3.6.0 via RStudio and plots were generated with the ggplot2 package. GLMM models used the glmer function and LMM models the lmer function, from the *lme4* package with *p* value approximation performed by the *lmerTest* package in R ^73,74^. Bayesian models were implemented using the *brms* package^70^. We ran a minimum of 2000 iterations over four MCMC chains, with a ‘warm-up’ period of 1000 iterations per chain leading to 4000 usable posterior samples, visual inspection of all MCMC results revealed satisfactory Rhat values (<1.01). In these analyses, an effect is seen as statistically significant if the credible interval does not contain zero with 95% certainty.

### Temporal interference (TI) stimulation

TI stimulation was delivered using a custom-made device as described in^14^. Two sinusoidal waveforms (at frequencies 2 kHz and 2.005 kHz) were supplied via a balanced pair of current sources that were driven in precisely opposite phase with a ground electrode carrying any imbalance currents (< 1%) from the paired current sources, preventing charging of the body relative to earth ground. Two pairs of stimulation electrodes (self-adhesive TENS, 1.5 cm x 1.5 cm with the corners cut to produce a rounded shape), were positioned on the participants’ heads using a conductive paste (Ten20, D.O. Weaver, Aurora, CO, USA) and held in place using medical tape (3M™ Micropore ™ medical tape). Electrode 1 (*e*1) and electrode 3 (*e*3) were positioned on the left hemisphere at the level of the nasion plane, *e*1 was positioned anterior to *e*3 (*e*1 at 50% of the subject’s half circumference minus 2.5 cm and *e*3 at 50% of the subject’s half circumference plus 2.5 cm, both counting from the nasion; such that the centres of the electrodes were 5 cm apart). Electrodes 2 and 4 (*e*2 and *e*4) were positioned on the right hemisphere at a plane just above the eyebrow, *e*2 was positioned anterior to *e*4 (*e*2 at 20% of the subject’s half circumference minus 1 cm and *e*4 at 70% of the subject’s half circumference plus 1 cm, both counting from the nasion). *e*1-*e*2 formed one electrode pair and *e*3-*e*4 the second electrode pair, **Fig. 1b**.

Stimulation was applied in two conditions: 1) TI stimulation directed to the mid left hippocampus: a current of 2 mA was applied to both electrode pairs, i.e., a current ratio 1:1 (‘TI 1:1’ condition); 2) TI stimulation steered to the anterior left hippocampus: a current of 1 mA was applied to the electrode pair *e1*-*e2* and a current of 3 mA was applied to the electrode pair *e3*-*e4*, i.e., a current ratio 1:3 (‘TI 1:3’ condition). In both conditions, the stimulation began with a 5 s ramp-up and ended with a 5 s ramp-down. Sham stimulation was equivalent to the TI 1:1 stimulation condition in the fMRI experiments, but the current was ramped down to zero immediately after it was ramped-up. In the behavioural experiment, sham stimulation was equivalent to the TI 1:3 condition, with an initial period of 30 s of stimulation followed by ramp-up and ramp-down periods before the first block of the face-name task and at the beginning and end of four consecutive blocks of the face-name task, see **Fig. S7**. The duration of stimulation was kept constant across participants for the fMRI experiment (96 s per stimulation condition), but durations varied between participants for the behavioural experiment where TI was applied throughout the face-name task blocks and responses were self-paced (stimulation during face-name task: mean ± SD, 34.5±3 min, range 29.37±40.35 min; total stimulation duration TI session: 44.53±3 min, range 39.37 – 50.35 min, which includes 10 min of stimulation before the face-name task, 5 min during rest and 5 min during a general alertness task, see **Fig. S7**).

The beginning and end of each stimulation block/period was controlled via an external trigger, sent from the computer running the experimental paradigm to the stimulator (triggers were sent from MATLAB using serial port commands). In the fMRI experiments, the start of each block was triggered by a signal from the MR scanner, this ensured that task and stimulation were synchronous to the scanner clock. The stimulator was placed outside the MR shielded room and the currents from the stimulator were delivered into the scanner room via RF filters, one placed in the operator room and another inside the scanner bore (NeuroConn GmbH, Ilmenau, Germany). The filter inside the MRI bore was connected to the stimulation electrodes via two MR compatible stimulation cables (NeuroConn GmbH, Ilmenau, Germany). Phantom and pilot experiments were initially conducted to ensure that the experimental setup did not introduce artifacts in the fMRI signal. Additionally, we estimated total signal-to-noise ratio (tSNR) in the fMRI signal in the brain regions underneath the electrodes on the left hemisphere and their contralateral equivalents (i.e., regions of interest, ROIs) to assess whether the presence of the electrodes had an effect on the quality of the signal. tSNR was calculated by dividing the mean of the signal over time by the standard deviation over the whole fMRI acquisition, **Fig. S8**.

### Brain stimulation procedure

Before each experiment, the participants’ sensation and comfort were tested. The participants were first exposed to low frequency stimulation followed by TI stimulation, for each electrode pair at a time, first *e*1-*e*2 followed by *e*3-*e*4. Stimulation started at 0.1 mA and increased in steps of 0.1 mA until participants reported any sensations associated to stimulation (i.e., pins and needles, burning, phosphenes, etc.) or until maximum intensities for the experimental protocol were reached (2 mA for *e*1-*e*2 and 3 mA for *e*3-*e*4). A detailed description of perceptual sensations and thresholds is provided in **Tables S18-S20**. At the end of the experiments, participants completed a questionnaire to assess possible side-effects of TI stimulation by rating from 0 (none) to 4 (severe) the intensity and duration of: pain, burning, warmth/heat, itchiness, pitching, metallic taste, fatigue, effect on performance, trouble concentrating, sleepiness/fatigue, headache, mood change, or any other side-effect perceived. A detailed description of side effects, their intensity and the number of incidences is reported in **Table S16** and **Table S17**. In the behavioural experiment we compared the strength ratings of the side effects using Wilcoxon signed-rank tests performed using the *coin* package^75^.

### Effectiveness of sham blinding

In the behavioural experiment, where we had separate active and sham sessions, we included an additional protocol to investigate the effectiveness of sham blinding by asking at 4 timepoints during each session the following questions: “Do you think you had stimulation” (yes/no) and “How confident are you?” (1 is not confident at all – 10 is extremely confident). Participants responded to each question by using an editable form in pdf format. Following^41^, we combined the two questions into a weighted score, whereby a “yes” answer was assigned a +1 value and “no” answer a value of -1, which were then multiplied by the confidence rating. We extracted the median and 95% confidence intervals for each time point and each stimulation condition using a smooth bootstrap technique^76^ implemented in the *kernelboot* package^77^. We used a Gaussian kernel and 10000 permutations for each probe point.

### MRI data acquisition

Scanning was performed in a 3T Siemens Verio (Siemens, Erlangen, Germany) at the Imperial College’s CIF, using a 32-channel head coil. Standard T1-weighted structural images were acquired using a magnetization-prepared rapid gradient-echo (MP-RAGE) sequence, 1 mm^3^ isotropic voxel, repetition time (TR) 2.3 s, echo time (TE) 2.98 ms, inversion time 900 ms, flip angle (FA) 9°, field of view 256 × 256 mm, 256 × 256 matrix, 160 slices, GRAPPA acceleration factor = 2. Field map scans were acquired to correct the echoplanar imaging (EPI) images for signal distortion (TR = 599 ms, TE = 7.65 ms, FA = 60°). FMRI images were obtained using a T2*-weighted gradient-echo EPI sequence, 3 mm^3^ isotropic voxel, TR 2 s, TE 30 ms, FA = 80°, field of view 192 × 192 × 105 mm, 35 slices, GRAPPA acceleration factor = 2. A total of 804 volumes were acquired on average (range: 592 – 1162), times varied depending on how long participants took on the recall stage of the task.

### Regions of interest

Regions of interest (ROIs) included: 1) the hippocampi, 2) their longitudinal parcellations; 3) regions corresponding to the AT-PM networks, and 4) regions corresponding to the cortical regions overlying the stimulated hippocampus, i.e., underneath and between the stimulating electrodes *e1* and *e3*.

Hippocampal masks were defined based on the segmentation of the whole hippocampi performed for each subject using the pipeline for automated hippocampal subfield segmentation in FreeSurfer (version 6.0.0,^78,79^). The hippocampal masks were normalised to MNI and split into thirds along the long axis of the hippocampus^80^ (posterior portion of the hippocampus: from Y= 40 to 30; mid-portion of the hippocampus: from Y= 29 to 19; anterior portion of the hippocampus: from Y= 18 to 4). The inverse normalization parameters were used to create subject specific parcellated ROIs and used in the subject space for fMRI analyses.

ROIs for the AT-PM networks were defined following^81^ using regions previously identified as belonging to distinct networks through resting state and functional connectivity analyses during associative memory encoding^82,83^. AT regions included the bilateral perirhinal cortex, amygdala, anterior fusiform gyrus, anterior inferior temporal cortex, and lateral orbitofrontal cortex. PM regions included the parahippocampal cortex, posterior cingulate cortex, precuneus and angular gyrus. The ROIs were obtained from probabilistic atlases thresholded at 50%, including a medial temporal lobe atlas (https://neurovault.org/collections/3731/; ^84^) for parahippocampal cortex and precuneus, and the Harvard-Oxford cortical and subcortical atlases for all other regions (**Fig. 4a**).

ROIs for the cortex overlying the stimulated hippocampus were defined for each subject using the anatomical T1 images. One ROI was placed underneath the anterior stimulating electrode *e1* (i.e., ROI Crtx Ant) and a second ROI was placed underneath the posterior stimulating electrode *e3* (i.e., ROI Crtx Post). The third ROI was placed in the middle between the electrodes (i.e., ROI Crtx Mid). All cortex ROIs were 10 mm spherical masks. See **Fig. 1d**.

We extracted two additional sets of ROIs for control measurements: the left amygdala (using a procedure analogous to the individual hippocampal ROIs), the left temporal lobe (excluding the hippocampus) and cortical ROIs in the right hemisphere (same axial plane as the left hemisphere ROIs). All ROIs were converted to the subject space for fMRI analyses.

### fMRI data pre-processing

Data were pre-processed using FMRIB Software Library (FSL version 6.0.1^85,86^. Functional data were pre-processed using the FMRI Expert Analysis Tool (FEAT), including motion correction using MCFLIRT^87^, distortion correction using fieldmap images prepared from fsl_prepare_fieldmap, slice-time correction using Slicetimer, smoothing with a 3D Gaussian kernel (8 mm full-width at half maximum, FWHM) and high-pass filtered at a cut-off of 0.008 Hz. Skull stripping was performed using FSL’s BET^88^. Head motion was estimated using FSL motion outliers through DVARS (the spatial root mean square of the data after temporal differencing)^89^. Criterion for excessive motion was DVARS > 0.5 in more than 20% of the volumes. In the fMRI experiment, one subject was excluded based on this. For the sample included in the analyses, mean DVARS and SD were 0.25 and 0.03 respectively. There was no difference in motion across stimulation conditions (F_(2,38)_=1.03, p=0.367).

### fMRI analysis

For each participant, pre-processed fMRI data was modelled using three different GLMs, two designed for univariate analyses and a third for assessing functional connectivity using generalised psychophysiological interaction (gPPI)^90^. In addition to the explanatory variables (EVs) of interest (described below), all GLMs included as nuisance regressors twenty-four motion parameters (six motion parameters - translation and rotation in three directions, the square of the six motion parameters and their temporal derivatives) and a regressor with volume outliers identified by DVARS to model out volumes (i.e., scrubbing) with extensive motion.

The first GLM was used to analyse univariate BOLD effects during encode and recall periods of the task and included 3 EVS for encode and 3 EVS for recall (one EV per stimulation condition and task stage) and their first temporal derivatives. Regressors were created by convolving a boxcar kernel with a canonical double-gamma hemodynamic response function.

The second GLM analysed univariate BOLD effects for correct and incorrect trials during encode and recall periods. This was possible without temporal jittering because we obtained a balanced distribution between correct and incorrect responses and the ordering of trials was randomised as a consequence of subject performance and pseudo-randomization of the stimuli presentation across participants^91^. This model included 12 EVs (one for correct and another for incorrect trials for encode and recall periods per stimulation condition), 3 EVs for the confidence intervals (one per stimulation condition) and their first temporal derivatives. Regressors were created by convolving a boxcar kernel with a canonical double-gamma hemodynamic response function.

The third set of GLMs, used to assess functional connectivity. We used a generalised psychophysiological interaction (gPPI) method to quantify the effective connectivity for the contrast correct > incorrect, using the Ant, Mid and Post regions of the left hippocampus as seeds and the AT and PM network as targets. The gPPI allowed us to quantify the directional connectivity between the seeds and targets while accounting for task-unrelated connectivity and task-related activity. The gPPI models included 25 EVs, describing physiological, psychological and PPI regressors. Physiological regressors were defined from the fMRI time-course extracted from seeds in the Ant, Mid and Post left hippocampus (see ROI definition). The psychological regressors included those modelled for the second GLM. For each model (one per seed), the physiological term and the psychological term were used to create the PPI interaction terms.

Using the output of the first GLM we assessed the fMRI BOLD signal to the encode and recall periods of the task (contrasted against the baseline), first for the sham condition (**Fig. 2**) and then for each stimulation condition (**Fig. 3**). Using the output of the second GLM, we measured BOLD response to correct and incorrect associations during the encode period of the task (contrasted against the baseline), first for the sham condition and then for all stimulation conditions (EVs 1-12). The contrast correct > incorrect was also used to extract connectivity values in the gPPI models described above.

### fMRI statistics

Whole brain BOLD activity at the group level was visualised by employing mixed effects analyses using FLAME 1^92,93^. Z statistical images were thresholded using Gaussian Random Fields based cluster inference with an initial cluster-forming threshold of Z>3.1 and a family-wise error (FWE) corrected cluster-extent threshold of p<0.05.

Statistical analyses of fMRI data were performed using the model estimates (in percent BOLD signal change) from the ROIs defined *a priori*. For each subject, we employed FSL Featquery tool to interrogate timeseries associated statistics, for each of the contrasts defined above, in the regions of interest in the subject space (see ROI specification). For analysis of the BOLD magnitude, we used the median % BOLD signal change (as the mean values were often not normally distributed) and for connectivity analysis we used the means.

ROI statistical analyses were conducted using R version 3.6.0 and plots were generated with the ggplot2 package. All mixed-effect models were fitted using the function lmer from the *lme4* package in R. ANOVA Type II Wald F tests with Kenward-Roger approximation for degrees of freedom or Type II Wald Chi-square tests were performed using the function Anova() for *p*-value approximation. Post hoc Tukey’s comparisons were made using the estimated marginal means from the *emmeans* package. The level of statistical significance was set at p<0.05 for all tests.

Voxelwise analyses within ROIs was performed using FSL’s randomise tool with 5,000 permutations and family-wise error correction for multiple comparisons using threshold-free cluster enhancement (TFCE). All statistical maps were family-wise corrected and thresholded at p < 0.05.

## Code and Data Availability

The code for the face-name task is available on Gitlab (https://gitlab.eps.surrey.ac.uk/nemo/facenametask). Data and key scripts are available on Gitlab (https://gitlab.eps.surrey.ac.uk/nemo/ti-paper). Group-level data used to generate the fMRI Fig.s are available in NeuroVault (https://neurovault.org/collections/11908/).

## ACKNOWLEDGEMENTS

I.R.V. is supported by the Biotechnology and Biological Sciences Research Council (BB/S008314/1). A.H. is supported by the UK Dementia Research Institute Care Research and Technology Centre and Biomedical Research Centre at Imperial College London. E.S.B was supported by Lisa Yang, John Doerr, HHMI, NIH R01MH117063, and Edward and Kay Poitras. N.G. was funded by the UK Dementia Research Institute (UK DRI)—an initiative funded by the Medical Research Council, Alzheimer’s Society and Alzheimer’s Research UK, Wellcome Trust fellowship (097443/Z/11/Z), Science & PINS Award for Neuromodulation, and NIHR IBRC Confident in Concept Award. A.W. is supported by the European Research Council (ERC) under the European Union’s Horizon 2020 research and innovation programme (716867). D.K is supported by the University of Surrey’s Vice-Chancellor Scholarship Award.

