## Supplemental Information for "Non-invasive temporal interference electrical stimulation of the human hippocampus"

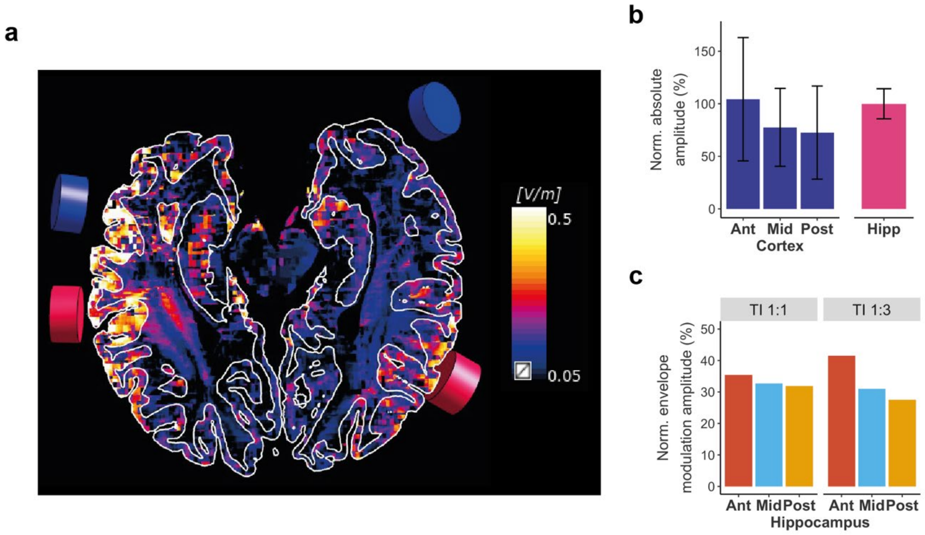

**Fig. S1: TI field distribution computed in the MIDA model.**
**a,** TI fields (i.e. envelope modulation amplitude) showed in an axial view parallel to the hippocampal longitudinal axis.
**b,** Absolute amplitude in regions of interest (ROIs) in the left (stimulated) hippocampus and its overlying cortex (see **Fig. 1d** for ROI locations and for the TI, i.e. envelope modulation amplitude); values are median±standard deviation (SD) normalised to the hippocampal value.
**c**, Envelope modulation amplitude in hippocampal ROIs (see **Fig. 2d** for ROI schematic) during TI 1:1 and TI 1:3 stimulations; ROI amplitudes were normalised to total hippocampal exposure.

**Table S1 | Induced potential in human cadaver; related to Fig. 1e.**
Induced potential for contacts in cortical and hippocampal tissue measured for the envelope modulation amplitude and absolute amplitudes, averaged across electrodes *a-c* (see **Fig. 1e**). Superficial contacts across the three intracranial electrodes were assigned to 'cortex' tissue (7 contacts, contacts 15-9, 12-33 mm depth) and deeper contacts to 'hippocampal' (8 contacts, contacts 8-1, 36.5-61 mm depth) tissue. Values represent the median ± standard deviation (SD). Statistics show paired samples t-test for envelope modulation amplitude and absolute amplitude between the two regions (i.e. cortex and hippocampus). Amplitudes in **Fig. 1e** have been normalised to hippocampal values.

| | **Cortex** | **Hippocampus** | **Statistics** |
|---|---|---|---|
| Envelope Modulation Amplitude mV | 0.35±0.38 | 1.22±0.34 | $t_{(4)} = -5.25$, $p = 0.006$ |
| Absolute Amplitude mV | 5.30±1.00 | 3.49±0.23 | $t_{(4)} = 7.05$, $p = 0.002$ |

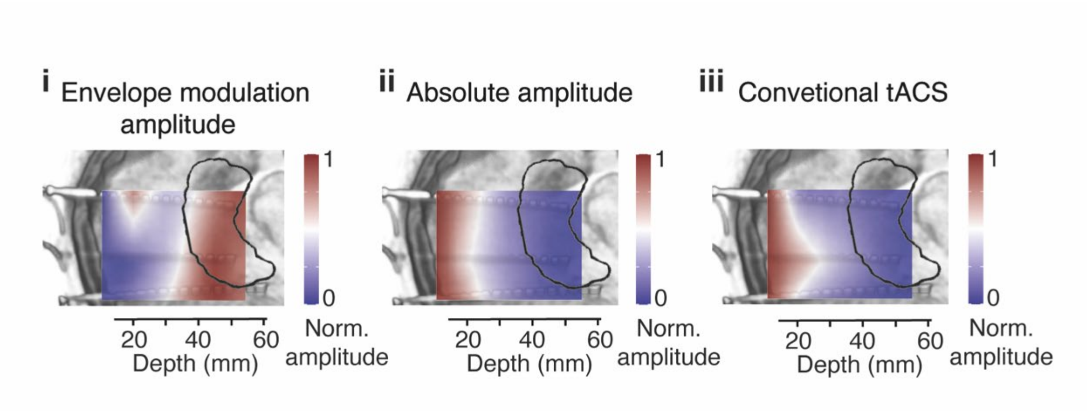

**Fig. S2: Measurements in human cadaver; related to Fig. 1.** Zoomed view of box region highlighted in **Fig. 1e** showing interpolated normalised amplitude maps of (**i**) envelope modulation amplitude map, (**ii**) absolute amplitude, (**iii**) absolute amplitude for conventional transcranial alternating current stimulation (tACS) at 5 Hz. Higher amplitudes are observed at the location of the hippocampus (black contour) for the envelope compared to absolute amplitude and conventional tACS.

**Table S2 | BOLD fMRI signal evoked by face-name memory task during sham (i.e., no) stimulation in the hippocampus; related to Fig. 2c.**
**S2.1.** One sample t-tests on the median BOLD signal (% signal change) extracted from individual hippocampal masks during encode and recall stages of the task, in the absence of stimulation, i.e. sham, condition. Statistical analyses were performed using one-sample t-tests, one-sided ("greater" than zero). P-values corrected using False Discover Rate (FDR). Shown are the estimate; t, t-statistic; P, P-value; P(FDR), P-value corrected using FDR. **S2.2**. Linear mixed effects model, with median BOLD (mdBOLD) signal as the dependent variable, independent factors for hemisphere (H: left, right) and task stage (TS: encode, recall), and random intercepts for participants (ID). Shown are the Analysis of Deviance Table (Type II Wald F tests with Kenward-Roger correction for degrees of freedom), generated by the Anova() function applied to the repeated measures analysis with mixed models fitted in R. [a]: Specification of the linear model fitted in the R language, Df, Degrees of freedom; Df.res, residual degrees of freedom; F, F-statistic; P, P-value; B, estimate; SE, standard error, t, t-statistic; N=20. Signif. codes: 0 '***' 0.001 '**' 0.01 '*' 0.05 '.' 0.1

| **S2.1 - One-sample t-test ("greater")** | | | | |
|---|---|---|---|---|
| **Task Stage** | **Hemisphere** | **t** | **P** | **P (FDR)** |
| Encode | Left | 3.6981 | 0.0008 | 0.0015 |
| | Right | 3.9215 | 0.0005 | 0.0015 |
| Recall | Left | -1.2812 | 0.8922 | 0.8922 |
| | Right | -0.2503 | 0.5975 | 0.7966 |

<table>
<tr><th colspan="5">S2.2 - Repeated measures analysis with mixed model</th></tr>
<tr><th>lmer(mdBOLD ~ TS*H + (1 | ID) [a]</th><th>Df</th><th>Df.res</th><th>F</th><th>P</th></tr>
<tr><td>Task Stage (TS)</td><td>1</td><td>57</td><td>20.4921</td><td>$3.097 \times 10^{-5}$ ***</td></tr>
<tr><td>Hemisphere (H)</td><td>1</td><td>57</td><td>0.5211</td><td>0.4733</td></tr>
<tr><td>Task Stage: Hemisphere</td><td>1</td><td>57</td><td>0.1328</td><td>0.7169</td></tr>
</table>

**Table S3 | BOLD fMRI signal evoked by face-name memory task during sham (i.e., no) stimulation in the segmented left hippocampus; related to Fig. 2e.**
LMM on the median BOLD signal (% signal change) extracted from individual segmentations of the left hippocampus during the encode stage of the task, in the absence of stimulation, i.e. sham, condition. Statistical analyses were performed using LMM, with median BOLD (mdBOLD) signal as the dependent variable, independent factors for ROI (Ant - anterior, Mid – mid, Post – posterior), and random intercepts for participants (ID). Shown are the Analysis of Deviance Table (Type II Wald F tests with Kenward-Roger correction for degrees of freedom), generated by the Anova() function applied to the repeated measures analysis with mixed models fitted in R, followed by the post-hoc contrasts for models with significant interactions. [a]: Specification of the linear model fitted in the R language, Df, Degrees of freedom; Df.res, residual degrees of freedom; F, F-statistic; P, P-value; B, estimate; SE, standard error, t, t-statistic N=20. Signif. codes: 0 '***' 0.001 '**' 0.01 '*' 0.05 '.' 0.1

| **Repeated measures analysis with mixed model** | | | | | |
|---|---|---|---|---|---|
| **lmer(mdBOLD ~ ROI + (1 \| ID) [a]** | **Df** | **Df.res** | **F** | **P** | |
| ROI | 2 | 38 | 8.7171 | 0.0007658 *** | |
| **Post-hoc contrasts** | | | | | |
| | **B** | **SE** | **df** | **t** | **P** |
| Ant - Mid | 0.1294 | 0.0322 | 38 | 4.024 | 0.0008 * |
| Ant - Post | 0.0957 | 0.0322 | 38 | 2.976 | 0.0137 * |
| Mid - Post | -0.0337 | 0.0322 | 38 | -1.048 | 0.5518 |

**Table S4 | BOLD fMRI signal evoked by face-name memory task during sham (i.e., no) stimulation in the segmented left hippocampus for the contrast correct > incorrect.**
**S4.1**. LMM for the median BOLD signal (% signal change) extracted from individual left hippocampal ROIs for fMRI Model 2, which differentiates between correct and incorrect responses during the encode stage of the task. Median BOLD (mdBOLD) signal is the dependent variable, independent factors for ROI (Ant - anterior, Mid – mid, Post – posterior) and response type (RspType: correct, incorrect), and random intercepts for participants (ID). Followed by estimated marginal means and standard error (SE).
**S4.2**. Same as S4.1 but for the right hippocampus.
Shown are the Analysis of Deviance Table (Type II Wald F tests with Kenward-Roger correction for degrees of freedom), generated by the Anova() function applied to the repeated measures analysis with mixed models fitted in R, followed by estimated marginal means for each factor for models with significant main effects. [a]: Specification of the linear model fitted in the R language, Df, Degrees of freedom; Df.res, residual degrees of freedom; F, F-statistic; P, P-value; Mean, estimated marginal mean; SE, standard error; N=20. Signif. codes: 0 '***' 0.001 '**' 0.01 '*' 0.05 '.' 0.1

| **S4.1 – Left Hippocampus** | | | | |
|---|---|---|---|---|
| **Repeated measures analysis with mixed model** | | | | |
| **lmer(mdBOLD ~ ROI*RspType + (1 \| ID) [a]** | **Df** | **Df.res** | **F** | **P** |
| ROI | 2 | 95 | 4.5808 | 0.012612 * |
| RspType | 1 | 95 | 11.0923 | 0.001235 ** |
| ROI: RspType | 2 | 95 | 0.9456 | 0.392065 |
| **Estimated Marginal Means** | | | | |
| **ROI** | **RspType** | **Mean** | | **SE** |
| Ant | Correct | 0.2748 | | 0.0547 |
| | Incorrect | 0.1151 | | 0.0547 |
| Mid | Correct | 0.1418 | | 0.0547 |

| | | | |
|---|---|---|---|
| | Incorrect | -0.0049 | 0.0547 |
| Post | Correct | 0.1236 | 0.0547 |
| | Incorrect | 0.0738 | 0.0547 |

| | | | | |
|---|---|---|---|---|
| **S4.2 – Right Hippocampus** | | | | |
| **Repeated measures analysis with mixed model** | | | | |
| **lmer(mdBOLD ~ ROI*RspType + (1 \| ID)** [a] | **Df** | **Df.res** | **F** | **P** |
| ROI | 2 | 95 | 0.1263 | 0.8815 |
| RspType | 1 | 95 | 1.3453 | 0.2490 |
| ROI: RspType | 2 | 95 | 0.1735 | 0.8410 |

**Table S5 | TI fields in the hippocampus; related to Fig. 2f.**
**S5.1**. Median and standard deviation (SD) for individualised electric field simulations based on participants' anatomical models, extracted from ROIs in the cortex and left hippocampus (see **Fig. 2d** for a schematic of the ROI locations). Shown are the envelope modulation and absolute amplitudes; N=16 (four subjects had to be excluded from the modelling since their electrodes were not visible in the MRI) for TI 1:1 and TI 1:3 stimulation conditions (2 mA current amplitude per electrode pair). **S5.2.** Statistical analyses on the normalised modulation envelope amplitudes for TI 1:1 and TI 1:3 stimulation conditions. Fields for each hippocampal region (**Fig. 2d)** were normalised to the whole hippocampus (analyses were conducted for the targeted left hippocampus, N=16). Linear mixed model (LMM) for each stimulation condition, followed by post-hoc pairwise comparisons. Models included the median normalised modulation envelope as the dependent variable (mdField), hippocampal regions (HippRg) as independent variable, and random intercepts for participants. Ant – anterior, Mid – middle, Post – posterior. Shown are the Analysis of Deviance Tables (Type II Wald F tests with Kenward-Roger correction for degrees of freedom), generated by the Anova() function applied to the linear mixed models fitted in R, followed by the post-hoc contrasts. [a]: Specification of the linear model fitted in the R language, Df, Degrees of freedom; Df.res, residual degrees of freedom; F, F-statistic; P: P-value; B, estimate; SE, standard error, t, t-statistic. Signif. codes: 0 '***' 0.001 '**' 0.01 '*' 0.05 '.' 0.1

| **S5.1 – Electric Field Simulation for Individualised Models** | | | |
|---|---|---|---|
| | | **Cortex** | **Hippocampus** |
| **TI 1:1** | Envelope Modulation Amplitude V/m | 0.23±0.18 | 0.40±0.05 |
| | Absolute Amplitude V/m | 0.46±0.25 | 0.47±0.05 |
| **TI 1:3** | Envelope Modulation Amplitude V/m | 0.19±0.15 | 0.24±0.03 |
| | Absolute Amplitude V/m | 0.45±0.26 | 0.47±0.05 |

| | | | | | |
|---|---|---|---|---|---|
| **S5.2 – Steering effect – Individualised Models** | | | | | |
| **Linear mixed model for TI 1:1** | | | | | |
| **lmer(mdField ~ HippRg + (1 \| ID)** [a] | **Df** | **Df.res** | **F** | **P** | |
| Hippocampal regions (HippRg) | 2 | 30 | 26.045 | $2.77 \times 10^{-7}$ *** | |
| **Post-hoc contrasts** | | | | | |
| | **B** | **SE** | **df** | **t** | **p** |
| Ant - Mid | -0.0487 | 0.00884 | 30 | -5.515 | <.0001 * |
| Ant - Post | 0.0113 | 0.00884 | 30 | 1.274 | 0.4203 |
| Mid - Post | 0.0600 | 0.00884 | 30 | 6.789 | <.0001 * |

| Linear mixed model for TI 1:3 | | | | | |
|---|---|---|---|---|---|
| **lmer(mdField ~ HippRg + (1 \| ID)** [a] | **Df** | **Df.res** | **F** | **P** | |
| Hippocampal regions (HippRg) | 2 | 30 | 359.62 | < 2.2 x$10^{-16}$ *** | |
| **Post-hoc contrasts** | | | | | |
| | **B** | **SE** | **df** | **t** | **p** |
| Ant - Mid | 0.0648 | 0.00552 | 30 | 11.741 | <.0001 * |
| Ant - Post | 0.1476 | 0.00552 | 30 | 26.752 | <.0001 * |
| Mid - Post | 0.0828 | 0.00552 | 30 | 15.011 | <.0001 * |

**Table S6 | BOLD signal in the hippocampus across stimulation conditions; related to Fig. 3b and Fig. 3g.**

**S6.1**. BOLD signal (% signal change) for the left, i.e. stimulated, hippocampus during encode and recall stages of the task, for the three stimulation conditions: sham, TI 1:1 and TI 1:3 (**Fig. 3b**). Statistical analyses were performed using a linear mixed effects model, with median BOLD (mdBOLD) signal as the dependent variable, independent factors for stimulation type (ST: sham, TI 1:1, TI 1:3) and task stage (TS: encode, recall), and random intercepts for participants (ID), N=20. **S6.2**. As per 6.1, but for the right, i.e. non-stimulated hippocampus; related to **Fig. 3g**; N=20**.**

| S6.1 – Left Hippocampus (stimulated hippocampus) | | | | | |
|---|---|---|---|---|---|
| **Repeated measures analysis with mixed model** | | | | | |
| **lmer(mdBOLD ~ ST*TS + (1 \| ID)** [a] | **Df** | **Df.res** | **F** | **P** | |
| Stimulation Type (ST) | 2 | 95 | 3.2224 | 0.04425 * | |
| Task Stage (TS) | 1 | 95 | 44.8436 | 1.492x$10^{-9}$ *** | |
| Stimulation Type:Task Stage | 2 | 95 | 2.9611 | 0.05656 . | |
| **Post-hoc contrasts** | | | | | |
| | **B** | **SE** | **df** | **t** | **P** |
| Task Stage = Encode | | | | | |
| Sham - TI 1:1 | 0.0124 | 0.042 | 95 | 0.296 | 0.9529 |
| Sham - TI 1:3 | 0.1317 | 0.042 | 95 | 3.133 | 0.0065 * |
| TI 1:1 - TI 1:3 | 0.1192 | 0.042 | 95 | 2.837 | 0.0153 * |
| Task Stage = Recall | | | | | |
| Sham - TI 1:1 | 0.0266 | 0.042 | 95 | 0.634 | 0.8020 |
| Sham - TI 1:3 | 0.0141 | 0.042 | 95 | 0.335 | 0.9399 |
| TI 1:1 - TI 1:3 | -0.0125 | 0.042 | 95 | -0.298 | 0.9522 |
| **S6.2 – Right Hippocampus** | | | | | |
| **Repeated measures analysis with mixed model** | | | | | |
| **lmer(mdBOLD ~ ST*TS + (1 \| ID)** [a] | **Df** | **Df.res** | **F** | **P** | |
| Stimulation Type (ST) | 2 | 95 | 0.7244 | 0.4873 | |
| Task Stage (TS) | 1 | 95 | 18.3715 | 4.359x$10^{-5}$ *** | |
| Stimulation Type:Task Stage | 2 | 95 | 2.0744 | 0.1313 | |

**Table S7 | BOLD signal in the segmented hippocampus across stimulation conditions; related to Fig. 3c.**

BOLD signal (% signal change) extracted from individual hippocampal segments during the encode stage of the task, for the three stimulation conditions: sham, TI 1:1 and TI 1:3. Statistical analyses were performed using a linear mixed effects model, with median BOLD (mdBOLD) signal as the dependent variable, independent factors for stimulation type (ST: sham, TI 1:1, TI 1:3) and ROI (Ant - anterior, Mid – mid, Post – posterior), and random intercepts for participants (ID), N=20. Shown are the Analysis of Deviance Table (Type II Wald F tests with Kenward-Roger correction for degrees of freedom), generated by the Anova() function applied to the repeated measures analysis with mixed models fitted in R, followed by estimated marginal means for each factor for models with significant main effects and difference to Sham condition. [a]: Specification of the linear model fitted in the R language, Df, Degrees of freedom; Df.res, residual degrees of freedom; F, F-statistic; P, P-value; Mean, estimated marginal mean; SE, standard error; N=20. Signif. codes: 0 '***' 0.001 '**' 0.01 '*' 0.05 '.' 0.1

| **Repeated measures analysis with mixed model** | | | | |
|---|---|---|---|---|
| **lmer(mdBOLD ~ ST*ROI + (1 \| ID)** [a] | **Df** | **Df.res** | **F** | **P** |
| Stimulation Type (ST) | 2 | 152 | 12.6459 | $8.313\times10^{-6}$ *** |
| ROI | 2 | 152 | 6.3506 | 0.002245 ** |
| Stimulation Type:ROI | 4 | 152 | 0.4639 | 0.762166 |
| **Estimated Marginal Means and Relative Differences to Sham** | | | | |
| | **Mean** | **SE** | **Mean – Sham** | |
| **ROI = Ant** | | | | |
| Sham | 0.2015 | 0.0501 | - | |
| TI 1:1 | 0.1931 | 0.0501 | 0.0084 | |
| TI 1:3 | 0.0247 | 0.0501 | 0.1770 | |
| **ROI = Mid** | | | | |
| Sham | 0.0721 | 0.0501 | - | |
| TI 1:1 | 0.0846 | 0.0501 | -0.0124 | |
| TI 1:3 | -0.0192 | 0.0501 | 0.0913 | |
| **ROI = Post** | | | | |
| Sham | 0.1058 | 0.0501 | - | |
| TI 1:1 | 0.0713 | 0.0501 | 0.0345 | |
| TI 1:3 | -0.0333 | 0.0501 | 0.1391 | |

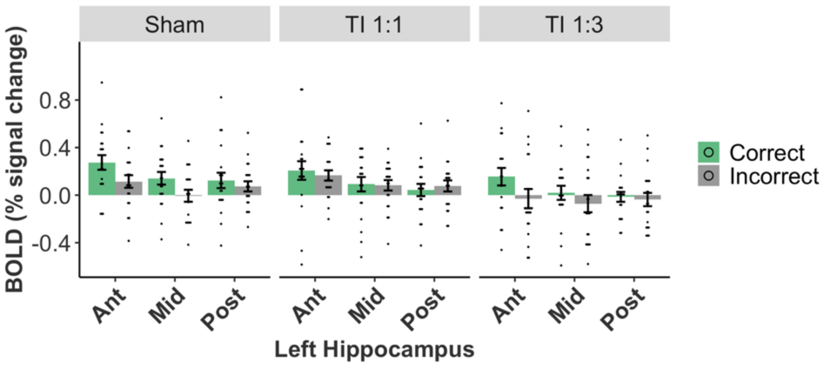

**Fig. S3: Comparison of group median change in BOLD signal between correct and incorrect encoded associations for each stimulation condition in the left hippocampus; related to Fig. 3e.** See **Table S4** and **Table S8** for full statistics.

**Table S8 | BOLD signal in the segmented hippocampus for correct and incorrect encoded associations; related to Fig. 2e.**

**S8.1**. Median BOLD signal (% signal change) extracted from individual left hippocampal ROIs during TI 1:3 stimulation for fMRI Model 2, which differentiates between correct and incorrect responses during the encode stage of the task. Statistical analyses were performed using a linear mixed effects model, with median BOLD (mdBOLD) signal as the dependent variable, independent factors for ROI (Ant - anterior, Mid – mid, Post – posterior) and response type (RspType: correct, incorrect), and random intercepts for participants (ID), N=20. **S8.2**. Same as S11.1 but for the TI 1:1 condition. Shown are the Analysis of Deviance Table (Type II Wald F tests with Kenward-Roger correction for degrees of freedom), generated by the Anova() function applied to the repeated measures analysis with mixed models fitted in R, followed by the post-hoc contrasts for models with significant interactions. [a]: Specification of the linear model fitted in the R language, Df, Degrees of freedom; Df.res, residual degrees of freedom; F, F-statistic; P, P-value; B, estimate; SE, standard error, t, t-statistic. Signif. codes: 0 '***' 0.001 '**' 0.01 '*' 0.05 '.' 0.1

| **S8.1 – Left Hippocampus Segmented – TI 1:3 condition** | | | | |
|---|---|---|---|---|
| **Repeated measures analysis with mixed model** | | | | |
| **lmer(mdBOLD ~ ROI*RspType + (1 \| ID)** [a] | **Df** | **Df.res** | **F** | **P** |
| ROI | 2 | 95 | 0.7125 | 0.49302 |
| RspType | 1 | 95 | 6.6161 | 0.01166 * |
| ROI: RspType | 2 | 95 | 1.4183 | 0.24722 |
| **Estimated Marginal Means** | | | | |
| **ROI** | **RspType** | **Mean** | **SE** | |
| Ant | Correct | 0.1123 | 0.0679 | |
| | Incorrect | -0.0672 | 0.0679 | |
| Mid | Correct | 0.0199 | 0.0679 | |
| | Incorrect | -0.0715 | 0.0679 | |
| Post | Correct | -0.0141 | 0.0679 | |
| | Incorrect | -0.0369 | 0.0679 | |
| **S8.2 – Left Hippocampus Segmented – TI 1:1 condition** | | | | |
| **Repeated measures analysis with mixed model** | | | | |
| **lmer(mdBOLD ~ ROI*RspType + (1 \| ID)** [a] | **Df** | **Df.res** | **F** | **P** |
| ROI | 2 | 95 | 4.7586 | 0.01073 * |
| RspType | 1 | 95 | 0.0273 | 0.86905 |
| ROI: RspType | 2 | 95 | 0.3992 | 0.67196 |
| **Estimated Marginal Means** | | | | |
| **ROI** | **RspType** | **Mean** | **SE** | |
| Ant | Correct | 0.2076 | 0.0553 | |
| | Incorrect | 0.1653 | 0.0553 | |
| Mid | Correct | 0.0921 | 0.0553 | |
| | Incorrect | 0.0828 | 0.0553 | |
| Post | Correct | 0.0434 | 0.0553 | |
| | Incorrect | 0.0776 | 0.0553 | |

**Table S9 | BOLD signal in cortical regions across stimulation conditions; related to Fig. 3h.**

**S9.1**. BOLD signal (% signal change) extracted from individual ROIs underneath and between the left hemisphere stimulation electrodes during encode stage of the task, for the three stimulation conditions: sham, TI 1:1 and TI 1:3. Statistical analyses were performed using a linear mixed effects model, with median BOLD (mdBOLD) signal as the dependent variable, independent factors for stimulation type (ST: sham, TI 1:1, TI 1:3) and ROI (Ant - anterior, Mid – mid, Post – posterior), and random intercepts for participants (ID), N=16. **S9.2.** As per 9.1, but for the left temporal lobe (excluding the hippocampus); N=20**.** Shown are the Analysis of Deviance Table (Type II Wald F tests with Kenward-Roger correction for degrees of freedom), generated by the Anova() function applied to the repeated measures analysis with mixed models fitted in R, followed by the post-hoc contrasts for models with significant interactions. [a]: Specification of the linear model fitted in the R language, Df, Degrees of freedom; Df.res, residual degrees of freedom; F, F-statistic; P, P-value; B, estimate; SE, standard error, t, t-statistic. Signif. codes:  0 '***' 0.001 '**' 0.01 '*' 0.05 '.' 0.1

| **S9.1 – Cortical ROIs** | | | | |
|---|---|---|---|---|
| **Repeated measures analysis with mixed model** | | | | |
| **lmer(mdBOLD ~ ST*ROI + (1 \| ID) [a]** | **Df** | **Df.res** | **F** | **P** |
| Stimulation Type (ST) | 2 | 120 | 2.4110 | 0.2604 |
| ROI | 2 | 120 | 10.1740 | $8.289 \times 10^{-5}$ *** |
| Stimulation Type:ROI | 4 | 120 | 0.1522 | 0.9617 |
| **S9.2 – Temporal Lobe** | | | | |
| **Repeated measures analysis with mixed model** | | | | |
| **lmer(mdBOLD ~ ST*TS + (1 \| ID) [a]** | **Df** | **Df.res** | **F** | **P** |
| Stimulation Type (ST) | 2 | 95 | 0.9704 | 0.3827 |
| Task Stage (TS) | 1 | 95 | 19.8443 | $2.294 \times 10^{-5}$ *** |
| Stimulation Type:Task Stage | 4 | 95 | 1.4511 | 0.2394 |

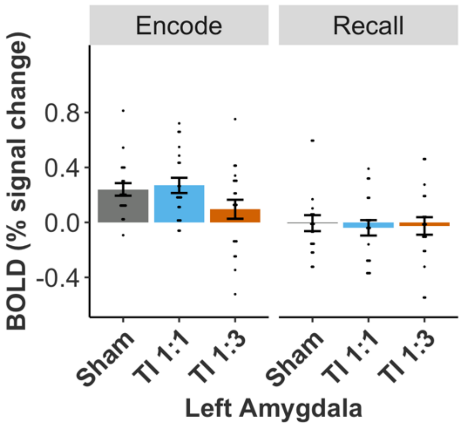

**Fig. S4: BOLD signal is not modulated by stimulation in the left amygdala.**
Group BOLD signal (% signal change) during encode and recall stages of the task across stimulation conditions (Sham, TI 1:1, TI 1:3). Statistical analyses were performed using a linear mixed effects model, with median BOLD signal as the dependent variable, independent factors for stimulation type (sham, TI 1:1, TI 1:3) and task stage (encode, recall), and random intercepts for participants, N=20. There is a main effect of task stage ($F_{(1,95)}$= 39.737, p= 9.08e-9), but no effect of stimulation type ($F_{(2,95)}$= 1.646, p= 0.198) and no interaction between task stage and stimulation type ($F_{(2,95)}$= 1.772, p= 0.176). Bar plots show mean and standard error (SE), black dots show individual participant data

**Table S10 | One sample t-tests for cortical Regions and left amygdala during sham stimulation.**
**S10.1.** One sample t-tests on the median BOLD signal (% signal change) extracted from individual cortical masks (see **Fig. 1d** for schematics of the location of the ROIs) during the encoding stage of the task, in the absence of stimulation, i.e. sham, condition. Statistical analyses were performed using one-sample t-tests, one-sided ("greater" than zero). P-values corrected using False Discover Rate (FDR). Shown are the estimate; t, t-statistic; P, P-value; P(FDR), P-value corrected using FDR; N=16. **S10.2**. Same as S10.1 but for the left amygdala; N=20. Signif. codes: 0 '***' 0.001 '**' 0.01 '*' 0.05 '.' 0.1

| **Cortical ROIs** | | | |
|---|---|---|---|
| **S10.1 - One-sample t-test ("greater")** | | | |
| **ROI** | **t** | **P** | **P (FDR)** |
| Crtx Ant | -0.8763 | 0.8027 | 0.8027 |
| Crtx Mid | 2.054 | 0.0289 * | 0.0434 |
| Crtx Post | 2.565 | 0.0108 * | 0.0323 |
| **Left Amygdala** | | | |
| **S10.2 - One-sample t-test ("greater")** | | | |
| **ROI** | **t** | **P** | **P (FDR)** |
| Left amygdala | 5.2502 | $2.280 \times 10^{-5}$ *** | $2.280 \times 10^{-5}$ |

**Table S11 | One sample t-tests for recall accuracy for face-name task performed during fMRI acquisition**
**S10.1.** One sample t-tests on the proportion of associations correctly recalled per stimulation condition. Statistical analyses were performed using one-sample t-tests, one-sided ("greater" than 0.2, which is the chance level, i.e. probability of selecting the target out of 5 possible responses). P-values corrected using False Discover Rate (FDR). Shown are the estimate; t, t-statistic; P, P-value; P(FDR), P-value corrected using FDR; N=20. Signif. codes: 0 '***' 0.001 '**' 0.01 '*' 0.05 '.' 0.1

| **Stimulation condition** | **t** | **P** | **P (FDR)** |
|---|---|---|---|
| Sham | 10.014 | $2.575 \times 10^{-9}$ | $2.575 \times 10^{-9}$ |
| TI 1:1 | 10.476 | $1.237 \times 10^{-9}$ | $1.856 \times 10^{-9}$ |
| TI 1:3 | 13.208 | $2.521 \times 10^{-11}$ | $7.563 \times 10^{-11}$ |

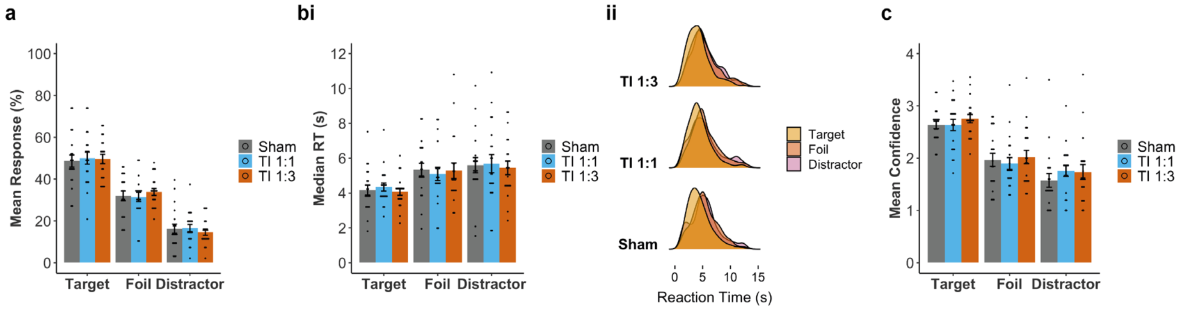

**Fig. S5: Comparison of participants' memory performances for face-name task, fMRI study.** Participant's performance across stimulation conditions, sham (grey), TI 1:1 (blue) and TI 1:3 (orange). **a**, Percentage mean response selection for each response category, showing a higher proportion of target selection compared to foils or distractors (probability of correct selection by chance was 0.2). **b, (i)** Median reaction time during recall and (**ii**) Recall time distribution across stimulation conditions for target (yellow), foil (orange) and distractor (pink) associations, showing faster reaction times for target selection. **c,** Mean confidence rating (from 1 low confidence to 4 high confidence) for each response category, showing higher confidence ratings for correct associations (i.e. target) compared to foils and distractors. See **Table S12** for full statistics.

**Table S12 | Memory performance statistics, fMRI study.**
Three main variables of interest were analysed to assess behavioural performance, i.e., response type – related to accuracy, reaction time for name selection and confidence level.
Shown are the statistical models applied to each behavioural metric (Model), the Analysis of Deviance Tables (Type II Wald chisquare tests), generated by the Anova() function applied to the models fitted in R. [a]: Specification of the model fitted in the R language, Df, Degrees of freedom; $\chi^2$, Chi-square; P, P-value; N=20 (ID). Signif. codes: 0 '***' 0.001 '**' 0.01 '*' 0.05 '.' 0.1

| **Accuracy** | | | |
|---|---|---|---|
| **Multinomial logistic regression** | | | |
| **nnet::multinom(formula= Response ~ ST)[a]** | **Df** | $\chi^2$ | **P** |
| Stimulation Type (ST) | 4 | 2.4288 | 0.6574 |
| **Binomial logistic regression** | | | |
| **glmer(Accuracy ~ ST + (1\| ID) + (1\|block), family = binomial(link = "logit"))[a]** | **Df** | $\chi^2$ | **P** |
| Stimulation Type (ST) | 2 | 0.0583 | 0.9713 |
| **Reaction Time** | | | |
| **Generalised mixed linear model** | | | |
| **glmer(RT ~ ST* Response + (1\|ID) + (1\|block), family = inverse.gaussian(link=identity)[a]** | **Df** | $\chi^2$ | **P** |
| Stimulation Type (ST) | 2 | 1.0652 | 0.5871 |
| Response (Target, Foil, Distractor) | 2 | 68.242 | $1.518 \times 10^{-15}$ *** |
| Stimulation Type:Response | 4 | 6.8863 | 0.1420 |
| **Generalised mixed linear model - binomial** | | | |
| **glmer(RT ~ ST*Accuracy + (1\|ID) + (1\|block), family = inverse.gaussian(link=identity))[a]** | **Df** | $\chi^2$ | **P** |
| Stimulation Type (ST) | 2 | 1.0700 | 0.58567 |
| Accuracy (Correct, Incorrect) | 1 | 69.212 | $< 2 \times 10^{-16}$ *** |

| | | | |
|---|---|---|---|
| Stimulation Type:Accuracy | 2 | 5.1867 | 0.07477 |
| **Confidence** | | | |
| **Cumulative Link Mixed Model** | | | |
| **clmm(Confidence ~ ST*Response + (1\|ID) + (1\|block), Hess = TRUE)[a]** | **Df** | $\chi^2$ | **P** |
| Stimulation Type (ST) | 2 | 10.43 | 0.005433 ** |
| Response (Target, Foil, Distractor) | 2 | 420.54 | $< 2 \times 10^{-16}$ *** |
| Stimulation Type:Response | 4 | 5.39 | 0.249476 |

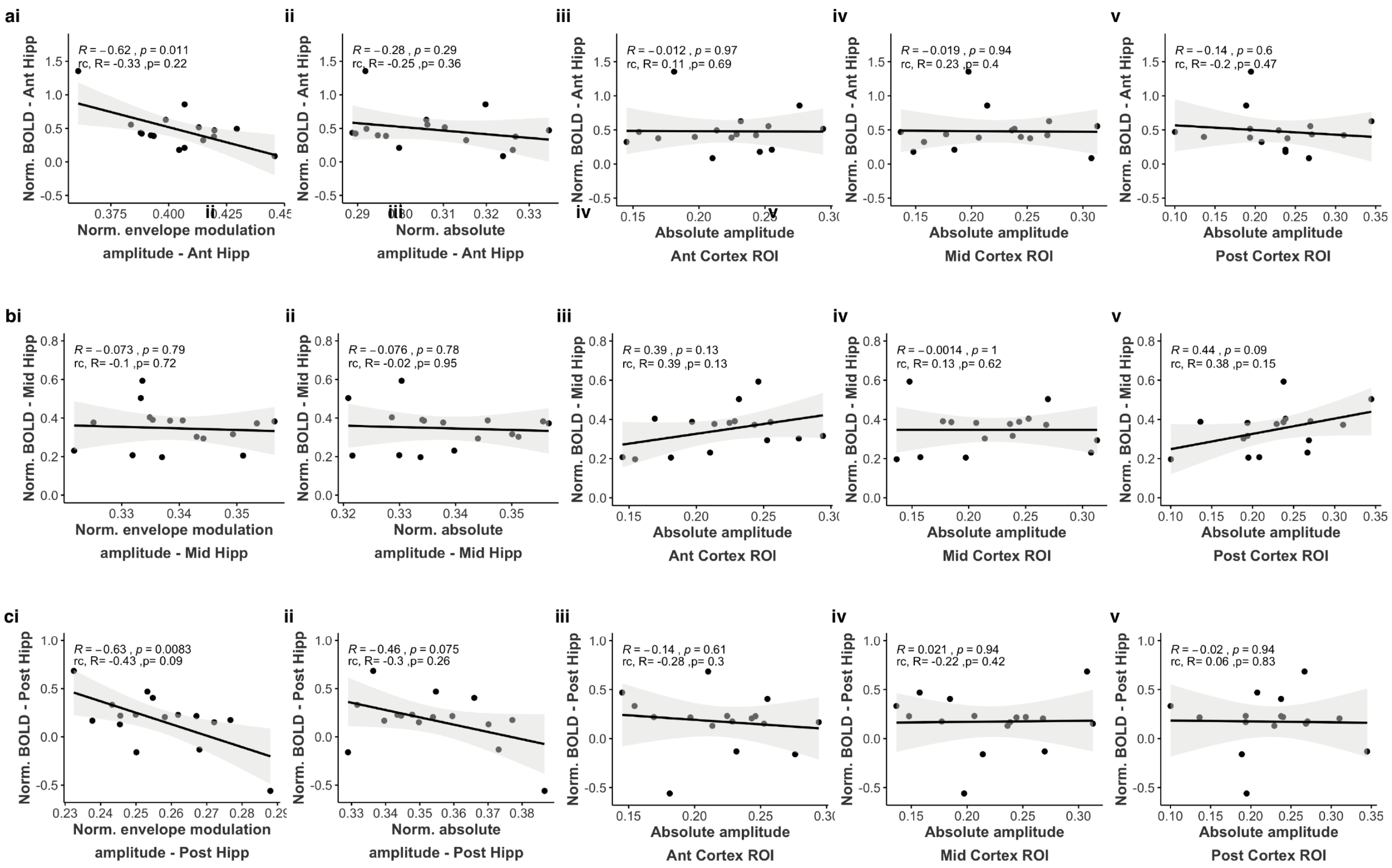

**Fig. S6:** Correlation between participants' evoked BOLD signal and their electric fields amplitudes for the TI 1:3 stimulation.

**a,** Correlation between BOLD signal from the Ant hippocampal and (i) envelope modulation amplitude in the Ant hippocampal region (ii) absolute amplitude in the Ant hippocampal region (expressed relative to total hippocampal exposure), (iii) absolute amplitude in the Ant overlying cortical region (iv) absolute amplitude in the Mid overlying cortical region (v) absolute amplitude in the Post overlying cortical region. The amplitude of the participants' evoked BOLD signal in the Ant left hippocampus during TI 1:3 stimulation was inversely correlated with the amplitude of the induced envelope modulation in this region (i, Pearson correlation but not robust correlation), but not with the absolute field amplitude in this region (ii) nor with the absolute field amplitude in the overlying cortical regions (iii-v). **b** and **c,** Same as (a), but using BOLD signal from the Mid (b) and Post (c) hippocampal region. Similar relationship between BOLD and TI fields observed for the Post region (c) as was observed in (a).

a-c. Hippocampal BOLD signal and field amplitudes in the hippocampus were normalized to total hippocampal exposure. For each plot, values at the top show the Pearson correlation coefficient (*R*), alongside p-values (*p*), regression lines (black line), confidence intervals of the correlation coefficient at 95% (shaded grey); also shown are the R and p-values calculated using robust correlation (rc), estimated using the *pbcor* function (bending factor = 0.2) from the WRS2 package [1]. N=16 subjects.

**Table S13 | Functional connectivity; related to Fig. 4.**

**S13.1**. Changes in functional connectivity for the sham condition during encode and recall stages of the task. The segmented regions on the left hippocampus were used as seeds (Ant, anterior; Mid, middle and Post, posterior) and nodes corresponding to the antero-temporal (AT) and posterior-medial (PM) networks as targets (see **Fig. 4a**). Functional connectivity was estimated using gPPI on the contrast correct > incorrect associations. Statistical analyses were performed using one-sample t-test, two sided, for the two task stages (encode, recall), three seeds (Ant, Mid, Post) and nodes bellowing to the AT or PM networks. The table shows the mean estimates (represented in **Fig. 3a**). t, T-statistic; P, P-value; P(FDR), P-value corrected with False Discovery Rate (FDR). N=20, **=p < 0.05, FDR-corrected; *=p < 0.05, uncorrected. **S13.2**. Functional connectivity values corresponding to the connectivity between each seed and target for each stimulation condition during the encode stage of the task (**Fig. 4c** – showing post-hoc contrasts). Statistical analyses were performed using a linear mixed effects model, with mean connectivity (mcon) as the dependent variable, independent factors for stimulation type (ST: sham, TI 1:1, TI 1:3), seed (S: Ant, anterior; Mid, middle; Post, posterior) and network (N: AT, PM), and random intercepts for participants (ID), and node from the AT or PM network. N = 20. Shown are the Analysis of Deviance Table (Type II Wald F tests with Kenward-Roger correction for degrees of freedom), generated by the Anova() function applied to the repeated measures analysis with mixed models fitted in R, followed by the post-hoc contrasts to assess the significant interactions. P value adjustment for post-hoc contrasts performed using Tukey's method (for a family of 3 estimates). [a]: Specification of the linear model fitted in the R language, Df, Degrees of freedom; Df.res, residual degrees of freedom; F, F-statistic; P: P-value; B, estimate; SE, standard error, t, t-statistic. Signif. codes: 0 '***' 0.001 '**' 0.01 '*' 0.05 '.' 0.1

| **S13.1 – Changes in functional connectivity for the sham condition** | | | | | | |
|---|---|---|---|---|---|---|
| **One sample t-test, two-sided** | | | | | | |
| **Task Stage** | **Seed** | **Target** | **estimate** | **t** | **P** | **P (FDR)** |
| Encode | Ant | AT | 0.1125 | 2.3221 | 0.0223 * | 0.1336 |
| Encode | Ant | PM | -0.0070 | -0.2546 | 0.7997 | 0.8724 |
| Encode | Mid | AT | 0.1445 | 3.1171 | 0.0024 * | 0.0287 * |
| Encode | Mid | PM | 0.0438 | 1.4547 | 0.1497 | 0.2756 |
| Encode | Post | AT | 0.0775 | 1.8025 | 0.0745 | 0.2756 |
| Encode | Post | PM | 0.0046 | 0.1550 | 0.8772 | 0.8772 |
| Recall | Ant | AT | 0.0482 | 1.6270 | 0.1069 | 0.2756 |
| Recall | Ant | PM | 0.0090 | 0.4087 | 0.6839 | 0.8207 |
| Recall | Mid | AT | 0.0538 | 1.4131 | 0.1608 | 0.2756 |
| Recall | Mid | PM | 0.0231 | 1.5282 | 0.1305 | 0.2756 |
| Recall | Post | AT | -0.0275 | -0.9292 | 0.3551 | 0.5326 |
| Recall | Post | PM | 0.0125 | 0.7563 | 0.4517 | 0.6023 |

| **S13.2 – Changes in connectivity across stimulation conditions** | | | | |
|---|---|---|---|---|
| **Repeated measures analysis with mixed model – Encode** | | | | |
| **lmer(mcon ~ ST*S*NT + (1 \| ID) + (1 \| node)** [a] | **Df** | **Df.res** | **F** | **P** |
| Stimulation Type (ST) | 2 | 1576 | 16.4165 | $8.784 \times 10^{-8}$ *** |
| Seed (S) | 2 | 1576 | 0.5299 | 0.58879 |
| Network (NT) | 1 | 1576 | 2.5686 | 0.15304 |
| Stimulation Type:Seed | 2 | 1576 | 2.6275 | 0.03305 * |
| Stimulation Type:Network | 2 | 1576 | 2.8940 | 0.05565 |
| Seed:Network | 4 | 1576 | 2.0604 | 0.12774 |
| Stimulation Type:Seed:Network | 4 | 1576 | 2.5382 | 0.03835 * |
| **Post-hoc contrasts** | | | | |

| | B | SE | df | t | P |
|---|---|---|---|---|---|
| Network = AT, seed = Ant | | | | | |
| Sham - TI 1:1 | 0.16982 | 0.0475 | 1576 | 3.577 | 0.0010 * |
| Sham - TI 1:3 | 0.02835 | 0.0475 | 1576 | 2.437 | 0.8216 |
| TI 1:1 - TI 1:3 | -0.14146 | 0.0475 | 1576 | 2.321 | 0.0082 * |
| Network = AT, seed = Mid | | | | | |
| Sham - TI 1:1 | 0.14638 | 0.0475 | 1576 | 3.084 | 0.0059 |
| Sham - TI 1:3 | 0.27135 | 0.0475 | 1576 | 5.717 | <.0001 * |
| TI 1:1 - TI 1:3 | 0.12497 | 0.0475 | 1576 | 2.633 | 0.0232 * |
| Network = AT, seed = Post | | | | | |
| Sham - TI 1:1 | 0.10521 | 0.0475 | 1576 | 2.216 | 0.0687 |
| Sham - TI 1:3 | 0.11532 | 0.0475 | 1576 | 2.429 | 0.0404 * |
| TI 1:1 - TI 1:3 | 0.01011 | 0.0475 | 1576 | 0.213 | 0.9753 |
| Network = PM, seed = Ant | | | | | |
| Sham - TI 1:1 | 0.04541 | 0.0531 | 1576 | 0.856 | 0.6684 |
| Sham - TI 1:3 | 0.04958 | 0.0531 | 1576 | 0.934 | 0.6187 |
| TI 1:1 - TI 1:3 | 0.00417 | 0.0531 | 1576 | 0.079 | 0.9966 |
| Network = PM, seed = Mid | | | | | |
| Sham - TI 1:1 | 0.07352 | 0.0531 | 1576 | 1.385 | 0.3488 |
| Sham - TI 1:3 | 0.04898 | 0.0531 | 1576 | 0.923 | 0.6259 |
| TI 1:1 - TI 1:3 | -0.02454 | 0.0531 | 1576 | -0.462 | 0.8888 |
| Network = PM, seed = Post | | | | | |
| Sham - TI 1:1 | 0.04004 | 0.0531 | 1576 | 0.754 | 0.7310 |
| Sham - TI 1:3 | 0.06538 | 0.0531 | 1576 | 1.232 | 0.4345 |
| TI 1:1 - TI 1:3 | 0.02534 | 0.0531 | 1576 | 0.478 | 0.8819 |

**Table S14 | Functional connectivity – comparison across stimulation conditions during recall period of the face-name memory task.**
**S14.1**. Functional connectivity values corresponding to the connectivity between each seed and target for each stimulation condition during the recall stage of the task. Statistical analyses were performed using a linear mixed effects model, with mean connectivity (mcon) as the dependent variable, independent factors for stimulation type (ST: sham, TI 1:1, TI 1:3), seed (S: Ant, anterior; Mid, middle; Post, posterior) and network (N: AT, PM), and random intercepts for participants (ID), and node from the AT or PM network. N = 20. **S14.2**. Follow-up LMM using only stimulation conditions as independent variable.
Shown are the Analysis of Deviance Table (Type II Wald F tests with Kenward-Roger correction for degrees of freedom), generated by the Anova() function applied to the repeated measures analysis with mixed models fitted in R, followed by the post-hoc contrasts to assess the significant interactions. P value adjustment for post-hoc contrasts performed using Tukey's method (for a family of 3 estimates). [a]: Specification of the linear model fitted in the R language, Df, Degrees of freedom; Df.res, residual degrees of freedom; F, F-statistic; P: P-value; B, estimate; SE, standard error, t, t-statistic. Signif. codes: 0 '***' 0.001 '**' 0.01 '*' 0.05 '.' 0.1

| **S14.1 – Changes in connectivity across stimulation conditions** | | | | |
|---|---|---|---|---|
| **Repeated measures analysis with mixed model – Recall** | | | | |
| **lmer(mcon ~ ST*S*NT + (1 \| ID) + (1 \| node) [a]** | **Df** | **Df.res** | **F** | **P** |
| Stimulation Type (ST) | 2 | 1576 | 8.3202 | 0.0002544 *** |
| Seed (S) | 2 | 1576 | 1.2781 | 0.2788424 |
| Network (NT) | 1 | 7 | 0.2214 | 0.6522743 |

| | | | | | |
|---|---|---|---|---|---|
| Stimulation Type:Seed | 2 | 1576 | 0.7631 | 0.5492473 | |
| Stimulation Type:Network | 2 | 1576 | 0.7193 | 0.4872524 | |
| Seed:Network | 4 | 1576 | 0.8136 | 0.4434303 | |
| Stimulation Type:Seed:Network | 4 | 1576 | 0.7121 | 0.5836283 | |
| **S14.2 – Changes in connectivity across stimulation conditions** | | | | | |
| **Repeated measures analysis with mixed model – Recall** | | | | | |
| **lmer(mcon ~ ST + (1 \| ID) [a]** | **Df** | **Df.res** | **F** | **P** | |
| Stimulation Type (ST) | 2 | 1598 | 8.2723 | 0.0002666 *** | |
| **Post-hoc contrasts** | | | | | |
| | **B** | **SE** | **df** | **t** | **P** |
| Sham - TI 1:1 | 0.05182 | 0.0145 | 1598 | 3.577 | 0.0010 * |
| Sham - TI 1:3 | 0.00169 | 0.0145 | 1598 | 0.117 | 0.9925 |
| TI 1:1 - TI 1:3 | -0.05013 | 0.0145 | 1598 | -3.463 | 0.0016 * |

**Table S15 | Memory performance statistics for face-name task; related to Fig. 5a.**
**S15.1**. Frequentist Analyses. Three main variables of interest were analysed to assess behavioural performance, i.e., response type – related to accuracy, reaction time for name selection and confidence level. Shown are the Analysis of Deviance Table (Type II Wald Chi-square or F tests with Kenward-Roger correction for degrees of freedom), generated by the Anova() function applied to the repeated measures analysis with mixed models fitted in R. [a]: Specification of the linear model fitted in the R language, Df, Degrees of freedom; Df.res, residual degrees of freedom; F, F-statistic; P: P-value; B, estimate; SE, standard error, t, t-statistic, Chisq – Chi-square. N=21. **S15.2.** Bayesian analysis. Results of Bayesian Regression Model for the effect of stimulation on accuracy (binomial distribution, i.e. correct and incorrect responses) estimated using the brms package in R. The specification of the model is shown in the R language, Est., the estimate of the mean marginal posterior distribution; Est. error, standard deviation of the estimate, CI, the 2.5% and 97.5% credible intervals centred on the mean, Post Dist. > 0, proportion of the posterior distribution greater than zero. N=21. Bayesian posterior density is plotted at the end of the table.

| **S15.1 – Frequentist Analyses** | | | | | |
|---|---|---|---|---|---|
| **Accuracy** | | | | | |
| **Multinomial logistic regression** | | | | | |
| **nnet::multinom(formula= Response ~ ST)[a]** | | **Df** | **$\chi^2$** | **P** | |
| Stimulation Type (ST) | | 2 | 6.353 | 0.04173 * | |
| **Post-hoc contrasts** | | | | | |
| | **B** | **SE** | **df** | **t** | **P** |
| Response = Target; Sham - TI 1:3 | -0.0282 | 0.0055 | 4 | -5.132 | 0.0068* |
| Response = Foil; Sham - TI 1:3 | 0.01968 | 0.0108 | 4 | 1.827 | 0.1418 |
| Response = Distractor; Sham - TI 1:3 | 0.00851 | 0.0087 | 4 | 0.978 | 0.3836 |
| **Binomial logistic regression** | | | | | |
| **glmer(Accuracy ~ ST + (1\| ID) + (1\|session) + (1\|block), family = binomial(link = "logit"))[a]** | | **Df** | **$\chi^2$** | **P** | |
| Stimulation Type (ST) | | 1 | 5.8567 | 0.01552 * | |
| **Reaction Time** | | | | | |
| **Generalised mixed linear model** | | | | | |

| glmer(RT ~ ST*Response + (1\|ID) + (1\|session) + (1\|block/trial), family = inverse.gaussian(link=identity), [a] | Df | $\chi^2$ | P |
|---|---|---|---|
| Stimulation Type (ST) | 1 | 3.0172 | 0.08239 |
| Response (Target, Foil, Distractor) | 2 | 424.71 | < 2 x$10^{-16}$ *** |
| Stimulation Type:Response | 2 | 2.8267 | 0.24333 |
| **Generalised mixed linear model - binomial** | | | |
| **glmer(RT ~ Accuracy*Response + (1\|ID) + (1\|session) + (1\|block/trial), family = inverse.gaussian(link=identity),** [a] | **Df** | $\chi^2$ | **P** |
| Stimulation Type (ST) | 1 | 2.9928 | 0.08364 |
| Response (Correct, Incorrect) | 1 | 425.47 | < 2 x$10^{-16}$ *** |
| Stimulation Type:Accuracy | 1 | 1.4945 | 0.22151 |
| **Confidence** | | | |
| **Cumulative Link Mixed Model** | | | |
| **clmm(Confidence ~ ST*Response + (1\|ID) + (1\|block), Hess = TRUE)**[a] | **Df** | $\chi^2$ | **P** |
| Stimulation Type (ST) | 1 | 0.35 | 0.5568 |
| Response (Target, Foil, Distractor) | 2 | 2006.1 | < 2 x$10^{-16}$ *** |
| Stimulation Type:Response | 2 | 0.42 | 0.8086 |

**S15.2 – Bayesian Analysis**

**Accuracy**

| Model | Predictor | Est. | Est. Error | CI 2.5% | CI 97.5% | Post Dist. > 0 |
|---|---|---|---|---|---|---|
| brm(Accuracy ~ Stimulation + (1\|ID) + (1\|session) + (1\|block), bernoulli(link = "logit"), prior = Prior_weak2.2, warmup = 2000, iter = 10000, chains = 4,cores = 4, control = list(adapt_delta = 0.999, max_treedepth = 17), seed = 1234)<br><br>ID = participant number<br>session = session number<br>block = block number | Intercept | 0.40 | 0.75 | -1.31 | 1.82 | - |
| | TI 1:3 | 0.12 | 0.05 | 0.02 | 0.22 | 99.21% |

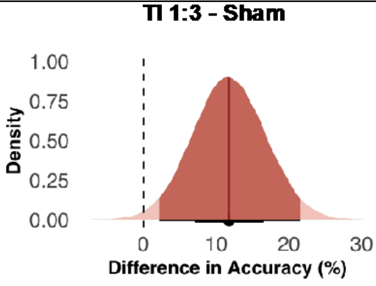

12%, 95% CI [2 22]

**Table S16 | Summary of post-stimulation side effects questionnaire for fMRI experiment**

Summary statistics for adverse effects questionnaire (adapted from [2]). Subjects rated each item between 1 (absent) and 4 (severe). Average Intensity and Range calculated over the whole cohort of participants.

| | Side effects questionnaire | | |
|---|---|---|---|
| | Average Intensity (SD) | Number of incidences > 1 (out of 21*) | Range |
| **Headache** | 1.75(0.5) | 4 | 1-2 |
| **Scalp pain** | 1.67(0.5) | 4 | 1-2 |
| **Burning** | 2(0) | 4 | 2 |
| **Warmth** | 2(0) | 4 | 2 |
| **Tingling** | 2.29(0.5) | 8 | 2-3 |
| **Prickling** | 2(0) | 3 | 2 |
| **Itching** | 1.67(0.5) | 4 | 1-2 |
| **Metallic taste** | 1(0) | 1 | 1 |
| **Sleepiness/Fatigue** | 2.57(0.9) | 8 | 1-4 |
| **Trouble concentrating** | 2.17(0.4) | 7 | 2-3 |
| **Effect on performance** | 2(0.6) | 6 | 1-3 |
| **Acute mood change** | 1(0) | 1 | 1 |
| **Other** | 1(0) | 1 | 1 |

* Initial sample size was 21. One participant was excluded from imaging analysis because of excessive movement. Information for all participants is included in the table. SD, standard deviation.

**Table S17 | Summary of post-stimulation side effects questionnaire for TI and Sham sessions for behavioural experiment**
Summary statistics for adverse effects questionnaire (adapted from [2]). Subjects rated each item between 1 (absent) and 4 (severe). Average Intensity and Range calculated over the whole cohort of participants.

| | **TI session** | | | **Sham session** | | | **Wilcoxon signed-rank tests** | |
|---|---|---|---|---|---|---|---|---|
| | Average Intensity (SD) | Number of incidences > 1 (out of 21) | Range | Average Intensity (SD) | Number of incidences > 1 (out of 21) | Range | Z | p |
| **Headache** | 1.2(0.5) | 3 | 1-3 | 1(0.2) | 1 | 1-2 | -1.063 | 0.288 |
| **Scalp pain** | 1.1(0.3) | 2 | 1-2 | 1.1(0.2) | 3 | 1-2 | 0.471 | 0.638 |
| **Burning** | 1.1(0.2) | 1 | 1-2 | 1(0) | 0 | 1 | -1 | 0.317 |
| **Warmth/Heat** | 1.1(0.3) | 2 | 1-2 | 1(0.2) | 1 | 1-2 | -0.592 | 0.554 |
| **Tingling** | 1.5(0.6) | 8 | 1-3 | 1.4(0.6) | 9 | 1-3 | 0 | 1 |
| **Itching** | 1.2(0.4) | 5 | 1-2 | 1(0) | 0 | 1 | -2.354 | 0.019* |
| **Metallic taste** | 1(0) | 0 | 1 | 1(0) | 0 | 1 | - | - |
| **Fatigue** | 1.6(0.7) | 10 | 1-3 | 1.6(0.8) | 9 | 1-3 | -0.112 | 0.911 |
| **Sleepiness** | 1.7(0.9) | 9 | 1-3 | 1.6(0.8) | 8 | 1-3 | -0.473 | 0.637 |
| **Acute mood change** | 1(0) | 0 | 1 | 1(0) | 0 | 1 | - | - |
| **Visual Sensation** | 1(0) | 0 | 1 | 1(0) | 0 | 1 | - | - |
| **Dizziness/ Nausea** | 1(0.2) | 1 | 1-2 | 1(0) | 0 | 1-2 | -1 | 0.317 |
| **Nervousness/ Anxiety** | 1(0.2) | 1 | 1-2 | 1(0) | 0 | 1-2 | -1 | 0.317 |
| **Discomfort/ Unpleasant** | 1.1(0.3) | 2 | 1-2 | 1(0) | 0 | 1-3 | -1.432 | 0.152 |
| **Other** | 1(0) | 0 | 1 | 1(0) | 0 | 1 | - | - |

**Table S18 | Perceptual sensations and threshold across participants (ID) for conventional transcranial alternating current stimulation (tACS) and TI stimulation for fMRI experiment**

Perceptual sensations reported by participants and thresholds (i.e. current intensity for which a perceptual sensation was first reported). Participants were exposed to short conventional tACS and TI stimulation during setup, immediately before entering the MRI scanner. TACS stimulation was always administered first, so participants would be aware of the possible sensations elicited by electrical stimulation. Current intensity was ramped in steps of 0.1 mA until a sensation was reported, starting from electrode pair *e*1-*e*2 and then moving to *e*3-*e*4.

| | **tACS (5 Hz)** | | | | **TI (CF = 2 and 2.005 kHz, $\Delta f$ = 5 Hz)** | | | |
|---|---|---|---|---|---|---|---|---|
| | ***e1 – e2*** | | ***e3 - e4*** | | ***e1 – e2*** | | ***e3 – e4*** | |
| **ID** | **Threshold (mA)** | **Sensation** | **Threshold (mA)** | **Sensation** | **Threshold (mA)** | **Sensation** | **Threshold (mA)** | **Sensation** |
| **1** | 0.3 | burning | 0.4 | light burning | none | none | 3 | stinging |
| **2** | 0.5 | prickling | 0.4 | stinging | none | none | none | none |
| **3** | 0.3 | tingling | 0.4 | tingling | none | none | none | none |
| **4*** | 0.3 | warmth | 0.5 | vibrating | none | none | none | none |
| **5** | 1 | vibration | 1.2 | tingling | none | none | none | none |
| **6** | 0.3 | prickling | 0.3 | tingling | none | none | none | none |
| **7** | 0.5 | prickling | 0.7 | itchy | none | none | none | none |
| **8** | 0.7 | prickling | 0.7 | warmth | none | none | none | none |
| **9** | 0.7 | tingling | 0.7 | tingling | none | none | none | none |
| **10** | 0.3 | prickling | 1.2 | tingling | none | none | none | none |
| **11** | 0.5 | prickling | 0.5 | warmth | none | none | none | none |
| **12** | 0.3 | prickling | 1.2 | tingling | none | none | none | none |
| **13** | 0.5 | stinging | 0.5 | tingling | none | none | none | none |
| **14** | 0.3 | tingling | 0.5 | tingling | none | none | none | none |
| **15** | 0.3 | tingling | 0.5 | tingling | 2 | tingling | 3 | tingling |
| **16** | 0.5 | prickling | 0.5 | prickling | none | none | none | none |
| **18** | 0.5 | tingling | 0.7 | tingling | none | none | none | none |
| **19** | 0.3 | tingling | 0.7 | tingling + warmth | none | none | 3 | pressure # |
| **20** | 0.3 | tingling | 0.5 | warmth | none | none | 3 | burning # |
| **21** | 0.5 | tingling | 0.5 | tingling + warmth | none | none | 3 | tingling # |
| **22** | 0.5 | stinging | 0.5 | stinging | none | none | none | none |

tACS – transcranial alternating current stimulation; TI – temporal interference stimulation; CF – carrier frequency; $\Delta f$ – delta frequency or modulated frequency; e – electrode. * Participant was excluded from imaging analysis because of excessive movement; # No sensations reported in the scanner.

**Table S19 | Perceptual sensations and threshold across participants (ID) for conventional transcranial alternating current stimulation (tACS) and TI stimulation – Session 1 behavioural experiment**

Perceptual sensations reported by participants and thresholds (i.e. current intensity for which a perceptual sensation was first reported). Participants were exposed to short conventional tACS and TI stimulation during setup, immediately before entering the MRI scanner. TACS stimulation was always administered first, so participants would be aware of the possible sensations elicited by electrical stimulation. Current intensity was ramped in steps of 0.1 mA until a sensation was reported, starting from electrode pair *e*1-*e*2 and then moving to *e*3-*e*4.

| | **tACS (5 Hz)** | | | | **TI (CF = 2 and 2.005 kHz, Δ*f* = 5 Hz)** | | | |
|---|---|---|---|---|---|---|---|---|
| | ***e*1 – *e*2** | | ***e*3 - *e*4** | | ***e*1 – *e*2** | | ***e*3 – *e*4** | |
| **ID** | **Threshold (mA)** | **Sensation** | **Threshold (mA)** | **Sensation** | **Threshold (mA)** | **Sensation** | **Threshold (mA)** | **Sensation** |
| **1** | 0.3 | pinprick | 0.3 | pinprick | 2 | pinprick | 2 | pinprick |
| **2** | 0.1 | stinging | 0.1 | stinging | 1.5 | stinging | 2 | stinging |
| **3** | 0.3 | pinprick | 0.3 | pinprick | 2 | tingling | 3 | tingling |
| **4** | 0.3 | tingling | 0.3 | tingling | 1.5 | vibrations | 1.5 | vibrations |
| **5** | 0.3 | tingling | 0.3 | tingling | none | none | none | none |
| **6** | 0.3 | pinprick | 0.7 | tingling | 1 | numbing | 2 | tingling |
| **7** | 0.5 | tingling | 0.3 | tingling | none | none | 2.5 | pressure |
| **8** | 0.5 | pinprick | 0.5 | pinprick | none | none | none | none |
| **9** | 0.3 | stinging | 0.5 | stinging | 2 | vibrations | 2 | vibrations |
| **10** | 0.1 | pinprick | 0.3 | pinprick | none | none | none | none |
| **11** | 0.1 | tingling | 0.3 | tingling | none | none | none | none |
| **12** | 0.5 | tingling | 0.5 | tingling | none | none | none | none |
| **13** | 0.7 | phosphenes | 0.5 | pulling | 1 | push/pull | 1 | push/pull |
| **14** | 0.3 | pinprick | 0.1 | pinprick | 1 | white noise | 1 | white noise |
| **15** | 0.3 | pinprick | 0.1 | heat | 2 | pinprick | 2 | pinprick |
| **16** | 0.1 | tingling | 0.3 | tingling | 2 | tingling | 2.5 | tingling |
| **17** | 0.3 | pinching | 0.3 | pinprick | none | none | 2 | hair prickle |
| **18** | 0.3 | pinprick | 0.3 | pinprick | none | none | none | none |
| **19** | 0.3 | pinprick | 0.5 | pinprick | 2 | tingling | none | none |
| **20** | 0.1 | pinprick | 0.1 | pinprick | 1.5 | pain | 1.5 | pain |
| **21** | 0.1 | pinprick | 0.5 | pinprick | 1 | laughter | 1 | laughter |

tACS – transcranial alternating current stimulation; TI – temporal interference stimulation; CF – carrier frequency; Δ*f* – delta frequency or modulated frequency; e – electrode.

**Table S20 | Perceptual sensations and threshold across participants (ID) for conventional transcranial alternating current stimulation (tACS) and TI stimulation – Session 2 behavioural experiment**

Perceptual sensations reported by participants and thresholds (i.e. current intensity for which a perceptual sensation was first reported). Participants were exposed to short conventional tACS and TI stimulation during setup, immediately before entering the MRI scanner. TACS stimulation was always administered first, so participants would be aware of the possible sensations elicited by electrical stimulation. Current intensity was ramped in steps of 0.1 mA until a sensation was reported, starting from electrode pair *e*1-*e*2 and then moving to *e*3-*e*4.

| | **tACS (5 Hz)** | | | | **TI (CF = 2 and 2.005 kHz, Δ*f* = 5 Hz)** | | | |
|---|---|---|---|---|---|---|---|---|
| | ***e*1 – *e*2** | | ***e*3 - *e*4** | | ***e*1 – *e*2** | | ***e*3 – *e*4** | |
| | **Threshold (mA)** | **Sensation** | **Threshold (mA)** | **Sensation** | **Threshold (mA)** | **Sensation** | **Threshold (mA)** | **Sensation** |
| **1** | 0.3 | tingling | 0.3 | tingling | 1.5 | vibrations | 2 | pinprick |
| **2** | 0.1 | tingling | 0.1 | tingling | 1.5 | stinging | 2 | stinging |
| **3** | 0.1 | pinprick | 0.3 | tingling | 2 | tickling | 3 | tingling |
| **4** | 0.3 | tingling | 0.3 | tingling | 1.5 | tingling | 1.5 | vibrations |
| **5** | 0.3 | tingling | *RM* | *RM* | 2 | twitching | none | none |
| **6** | 0.3 | tingling | 0.3 | tingling | 2 | tingling | 2 | tingling |
| **7** | 0.5 | tingling | 0.3 | tingling | none | none | 2.5 | pressure |
| **8** | 0.3 | pinprick | 0.3 | pinprick | none | none | none | none |
| **9** | 0.3 | tingling | 0.3 | tingling | none | none | 2 | vibrations |
| **10** | 0.1 | pinprick | 0.5 | pinprick | 2 | none | none | none |
| **11** | 0.1 | tingling | 0.3 | tingling | none | none | none | none |
| **12** | 0.3 | tingling | 0.3 | mild tingling | 2 | none | none | none |
| **13** | 0.7 | piercing | 0.9 | none | 2 | slight pulling in occipital region | 1 | push/pull |
| **14** | 0.1 | pinprick | 0.5 | faster pinpricks | 1 | white noise | 1 | white noise |
| **15** | 0.1 | pinprick | 0.3 | pinprick | 1.5 | pinprick | 2 | pinprick |
| **16** | 0.3 | touch | 0.7 | sting | 2 | tingling | 2.5 | tingling |
| **17** | 0.5 | pinprick/small needle | 0.5 | pinprick/ small needle | 1 | light tapping | 2 | hair prickle |
| **18** | 0.1 | tingling | 0.1 | tingling | none | none | none | none |
| **19** | 0.3 | tingling | 0.5 | slight tingling | 1 | hair moving | none | none |
| **20** | 0.1 | tingling | 0.1 | tingling | 1.5 | itchy | 1.5 | pain |
| **21** | 0.1 | pinprick | 0.5 | pinprick | 1.5 | headache | 1 | laughter |

tACS – transcranial alternating current stimulation; TI – temporal interference stimulation; CF – carrier frequency; Δ*f* – delta frequency or modulated frequency; e – electrode. RM – record missing

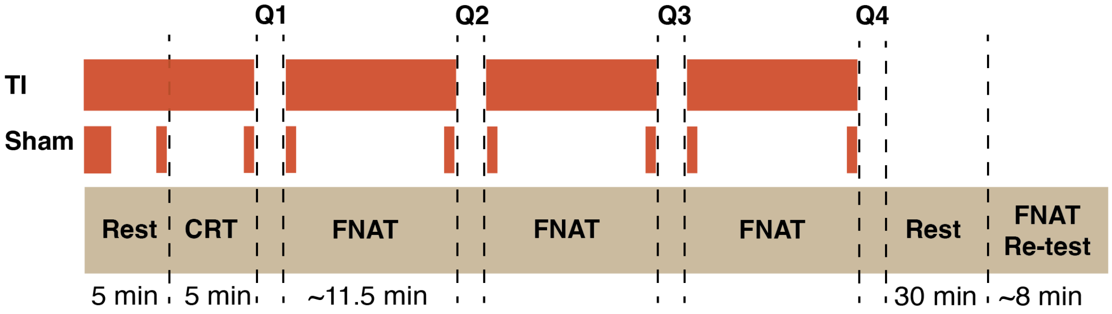

**Fig. S7. Experimental design for behavioural experiment.**
Red horizontal bars indicate stimulation periods; Q1-4, stimulation blindness questionnaire 1 to 4. FNAT, Face-Name Task; CRT, Choice-Reaction Time Task. The CRT is a simple 2-alternative force choice paradigm commonly used for testing general alertness and motor speed. Participants were presented with a central right or left pointing arrows and asked to press a right or left button matching the direction of the arrow, as described in[3]. The task consisted of 136 ± 1 trials, 1 ± 0.2 s inter stimulus interval (ISI, randomly drawn for each trial from a normal distribution with a mean of 1 s and variance of 0.2 s). Accuracy and reaction times were similar between stimulation conditions. There was no difference in accuracy between conditions (LMM: $\chi^2(1) = 0.179$, p= 0.672, accuracy mean±SD, sham: 99.7±5.4%; TI: 99.6±6%) or median reaction times (LMM: $\chi^2(1)= 3.507$, p= 0.0611, reaction time for correct trials median±SD, sham 0.394± 0.073 s; TI: 0.398± 0.076 s).

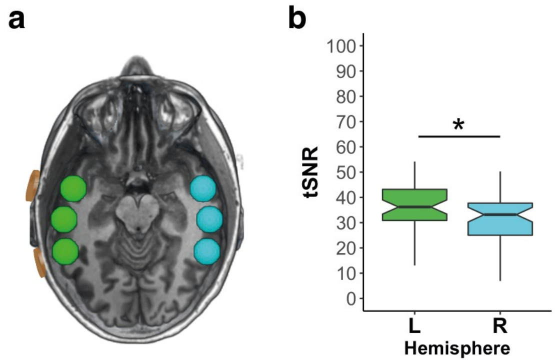

**Fig. S8: Total Signal to Noise Ratio (tSNR) in cortical ROIs.**
**a,** To assess whether the presence of electrodes on the scalp affected the quality of the fMRI images, we estimated total signal-to-noise ratio (tSNR) in the brain regions underneath and between the electrodes on the left hemisphere (green) and their contralateral equivalents (cyan, right hemisphere ROIs). The location of the stimulation electrodes on the left hemisphere are represented in orange for an example participant. **b,** Group tSNR for the left (L) and right (R) hemisphere ROIs (Median BOLD signal; centre line, median; box limits, upper and lower quartiles; lines, 1.5 x interquartile range; *p<0.05). tSNR was calculated by dividing the mean of the signal over time by the standard deviation over the whole fMRI acquisition for Experiment 1 (Face-name task, for N=16 where electrodes were clearly visible on T1 images). A linear mixed model (LMM) with tSNR as the dependent variable and hemisphere as the independent variable, and random intercepts for participants and ROI indicated a main effect of hemisphere ($F_{(1,77)}$= 17.175, p= 8.675e$^{-05}$) explained by higher tSNR in the left compared to the right hemisphere (post-hoc contrasts: $t_{(77)}$= 4.144, p=0.0001). This indicates that there was no reduction in tSNR underneath and between the electrodes in the left hemisphere, and in fact tSNR was higher in the left compared to the right hemisphere. These results indicate that

our electrodes and stimulation equipment did not introduce the patterns of noise in the MR signal identified in some studies conducting simultaneous brain stimulation and fMRI, which are typically characterised by a reduction of tSNR underneath the stimulation electrodes [4].
